# RNA- and DNA-binding proteins generally exhibit direct transfer of polynucleotides: Implications for target site search

**DOI:** 10.1101/2022.11.30.518605

**Authors:** Wayne O. Hemphill, Calvin K. Voong, Regan Fenske, James A. Goodrich, Thomas R. Cech

**Author notes:** **Author contributions:** W.O.H., C.K.V., J.A.G. and T.R.C. designed research; W.O.H., C.K.V. and R.F. performed research; W.O.H. and R.F. analyzed data; and W.O.H. and T.R.C. wrote the paper. **Competing Interest Statement:** T.R.C declares consulting status for Storm Therapeutics, Eikon Therapeutics, and SomaLogic. W.O.H. declares the filing of U.S. Provisional Application No. 62/706,167 Trex1 Inhibitors and Uses Thereof. The authors have no other competing interests to declare.

## Abstract

We previously demonstrated that the PRC2 chromatin-modifying enzyme exhibits the ability to directly transfer between RNA and DNA without a free-enzyme intermediate state. Simulations suggested that such a direct transfer mechanism may be generally necessary for RNA to recruit proteins to chromatin, but the prevalence of direct transfer capability is unknown. Herein, we used fluorescence polarization assays and observed direct transfer for several well-characterized nucleic acid-binding proteins: three-prime repair exonuclease 1 (TREX1), heterogeneous nuclear ribonucleoprotein U, Fem-3-binding factor 2, and MS2 bacteriophage coat protein. For TREX1, the direct transfer mechanism was additionally interrogated by single molecule assays, and the data suggest that direct transfer occurs through an unstable ternary intermediate with partially associated ligands. Generally, direct transfer could allow many DNA- and RNA-binding proteins to conduct a one-dimensional search for their target sites. Furthermore, presumably long-lived protein-polynucleotide complexes might instead be readily replaced by other protein-polynucleotide complexes *in vivo*.

**Significance:** Classically, the lifetime of a protein-ligand complex is presumed to be an intrinsic property, unaffected by competitor molecules in free solution. By contrast, a few oligomeric nucleic acid binding proteins have been observed to exchange competing ligands in their binding sites, and consequently their lifetimes decrease with competitor concentration. Our findings indicate that this “direct transfer” is a more general property of nucleic acid binding proteins. This suggests that many DNA- and RNA-binding proteins can reduce the dimensionality of their search for their target sites by intramolecular direct transfer to nucleosome DNA, instead of relying entirely on three-dimensional diffusion, and it suggests that their mean complex lifetimes *in vivo* can be regulated by the concentration of free ligand molecules.

## Introduction

A few well-characterized oligomeric proteins have been shown to exchange ligand species through highly unstable ternary complex intermediates (i.e., “direct transfer”) (1–5). In concurrent studies (companion manuscript), we demonstrated that this phenomenon of direct transfer occurs between RNA and DNA for the chromatin-modifying enzyme Polycomb Repressive Complex 2 (PRC2) (6); this phenomenon could allow mutually antagonistic RNA and DNA binding to both positively and negatively regulate PRC2 activity depending on the transcriptional environment, contrary to the classic assumptions. The prevalence of this direct transfer phenomenon among RNA- and DNA-binding proteins is not known. Thus, a robust study of direct transfer with various nucleic acid binding proteins (NBPs) was warranted.

NBPs with well-characterized biochemistry span a wide range of biological functions. Heterogeneous nuclear ribonucleoprotein U (hnRNP-U) is an RNA- and DNA-binding protein proposed to regulate chromatin structure and pre-mRNA processing (7). Recent work has also demonstrated that hnRNP-U has specificity for G-quadruplex (G4) RNA (8), much like PRC2 (9). Three-prime repair exonuclease 1 (TREX1) (10, 11) is a 3’→5’ exonuclease (12) that degrades DNA to prevent aberrant nucleic acid sensing (13) and the resulting harsh autoimmunity (14). In recent years, interest in TREX1 research has risen due to its emerging potential as a cancer immunotherapy target (15), and TREX1 activity on ss-versus dsDNA, the purpose of its homodimer structure, and the source of TREX1’s DNA substrates *in vivo* remain areas of active interest (16, 17). Fem-3-binding factor 2 (FBF-2) is a Pumilio Factor (PUF) family sequence-specific RNA-binding protein that binds the 3’-untranslated regions of mRNAs to inhibit expression of the protein machinery necessary for meiotic entry during *C. elegans* germline development (18–20). MS2 bacteriophage coat protein (MS2-CP) is a capsid component necessary for viral particle assembly of this *E. coli* bacteriophage, and it also negatively regulates MS2 replicase expression (21, 22). MS2-CP exhibits sequence-specific hairpin RNA binding, which is necessary for these functions (23–26).

Here we use a variety of biophysical assays to interrogate the prevalence, mechanism, and biophysical requisites of direct transfer among these NBPs. Our findings indicate that direct transfer occurs when ligands compete for shared protein contacts and partially associate to form an unstable ternary complex intermediate. They imply that the polydentate nature of protein-polynucleotide interactions makes direct transfer a common feature of NBPs. This supports direct transfer being a feature of many RNA-binding chromatin-associated proteins that may be generally required for their tunable regulation (companion manuscript), and there are also implications for NBPs in general. Notably, prior work has suggested that direct transfer could allow for protein movement along DNA via “looping” (27–29), which would allow many NBPs to benefit from a one-dimensional search for their gene targets after initial chromatin association. Similarly, RNA-binding proteins could be transferred intramolecularly and intermolecularly in search for optimal binding sites. In addition, the apparent mechanistic relation of direct transfer to the phenomenon of “facilitated dissociation” (FD) (30) has the transitive implication that NBP complexes *in vivo* can be prematurely displaced by other binders in the reaction environment, instead of being rate-limited by intrinsic dissociation of the complex. These implications substantially expand the possibilities for how NBPs find their respective binding partners and regulate their biological activity.

## Results

### Diverse Protein-Polynucleotide Interactions Exhibit Direct Transfer Kinetics

The ability of a protein to directly exchange one polynucleotide for another, without a free protein intermediate state, has important implications for mechanisms of proteins finding their binding sites within the cell and the lifetimes of these complexes *in vivo*. To assess the generality of this mechanism, we selected a variety of well-characterized protein-nucleic acid interactions, including hnRNP-U binding to G4 RNA (8), TREX1 binding to short ssDNA (10, 11), FBF-2 binding to a short ssRNA sequence (18, 19), and MS2-CP binding to its RNA hairpin motif (23, 24). We also tested streptavidin binding to biotin (31). These five protein-ligand systems were chosen for their collective diversity in structure, type of bonding between protein and ligand, ligand specificity, binding surface area, affinity, and stoichiometry.

We used purified recombinant protein and fluorescence polarization (FP) experiments to determine K_d_^app^ for these interactions (Table 1). Our binding affinity results were consistent with prior reports for TREX1 (32), hnRNP-U (8), MS2-CP (25), FBF-2 (19), and streptavidin (31). Then, we performed FP-based competitive dissociation (FPCD) experiments (Supp. Fig. 1) with prey (fluorophore-labeled) and decoy (unlabeled) ligands of the same identity to determine which protein-ligand interactions, if any, had an apparent dissociation rate (k_off_^obs^) that was dependent on the concentration of the decoy ligand, versus plateauing at high decoy concentrations. In other words, whether the dissociation rate was best described by one versus two rate constants: one for classic dissociation (k_-1_) and an optional second for direct transfer (k_θ_). Unexpectedly, our results (Fig. 1) revealed decoy concentration-dependent dissociation kinetics for every NBP (i.e., direct transfer), with their second-order rate constants for direct transfer spanning a 40-fold range. Only the streptavidin-biotin interaction had an apparent dissociation rate that plateaued at high decoy concentrations (i.e., classic competition). The data for the NBPs were consistent with a direct transfer mechanism, where competing ligand directly attacks the protein-nucleic acid complex and facilitates displacement of the existing ligand.

**Table 1.**
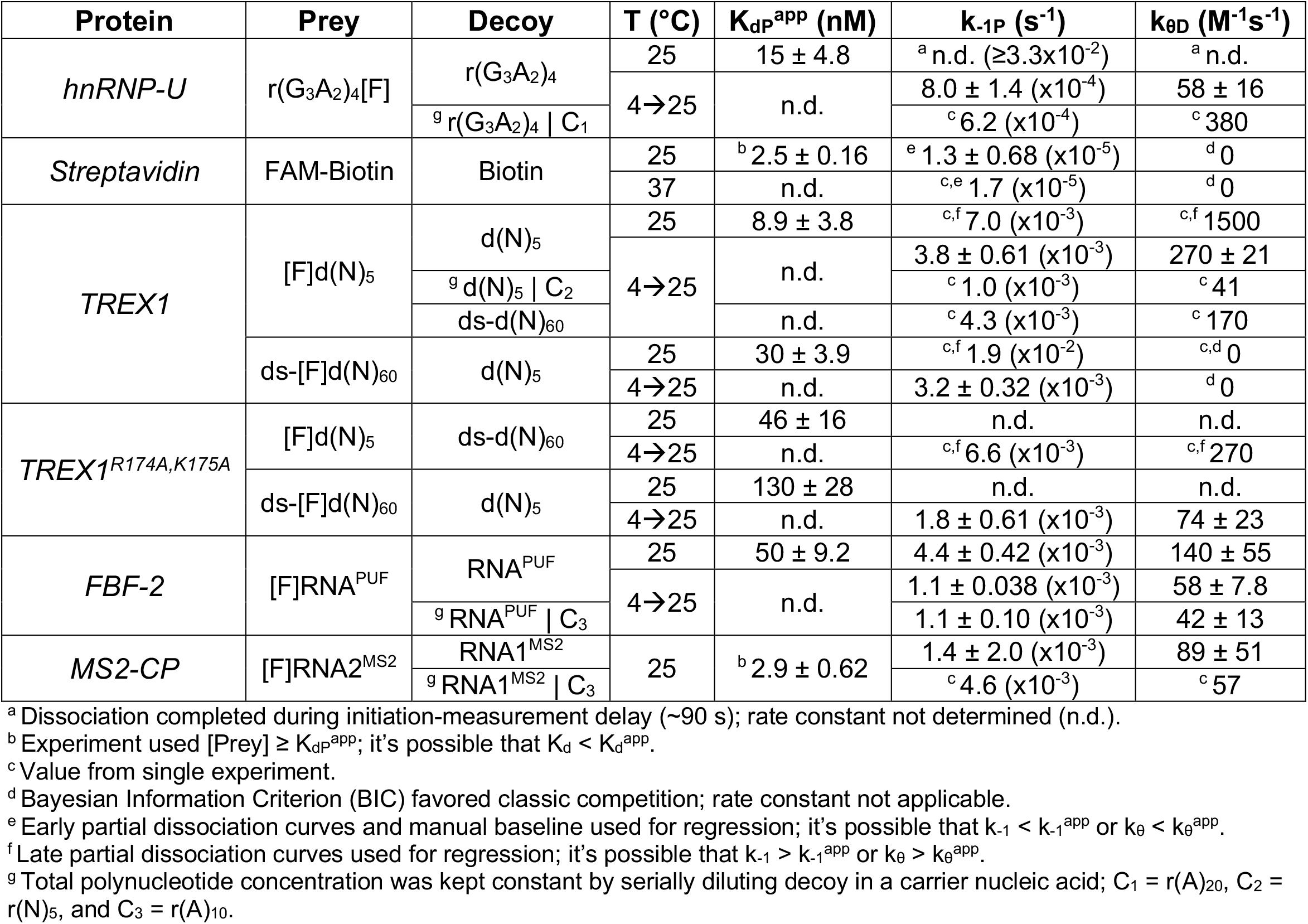
Rate Constants for Protein-Ligand Interactions. Fluorescence polarization-based methodology (Supp. Fig. 1) was used to determine the apparent equilibrium dissociation constants (K_dP_^app^), intrinsic dissociation rate constants (k_-1P_), and direct transfer rate constants (k_θD_) for several protein-ligand interactions. Values indicate mean ± SD for at least three independent experiments.

**Fig. 1.**
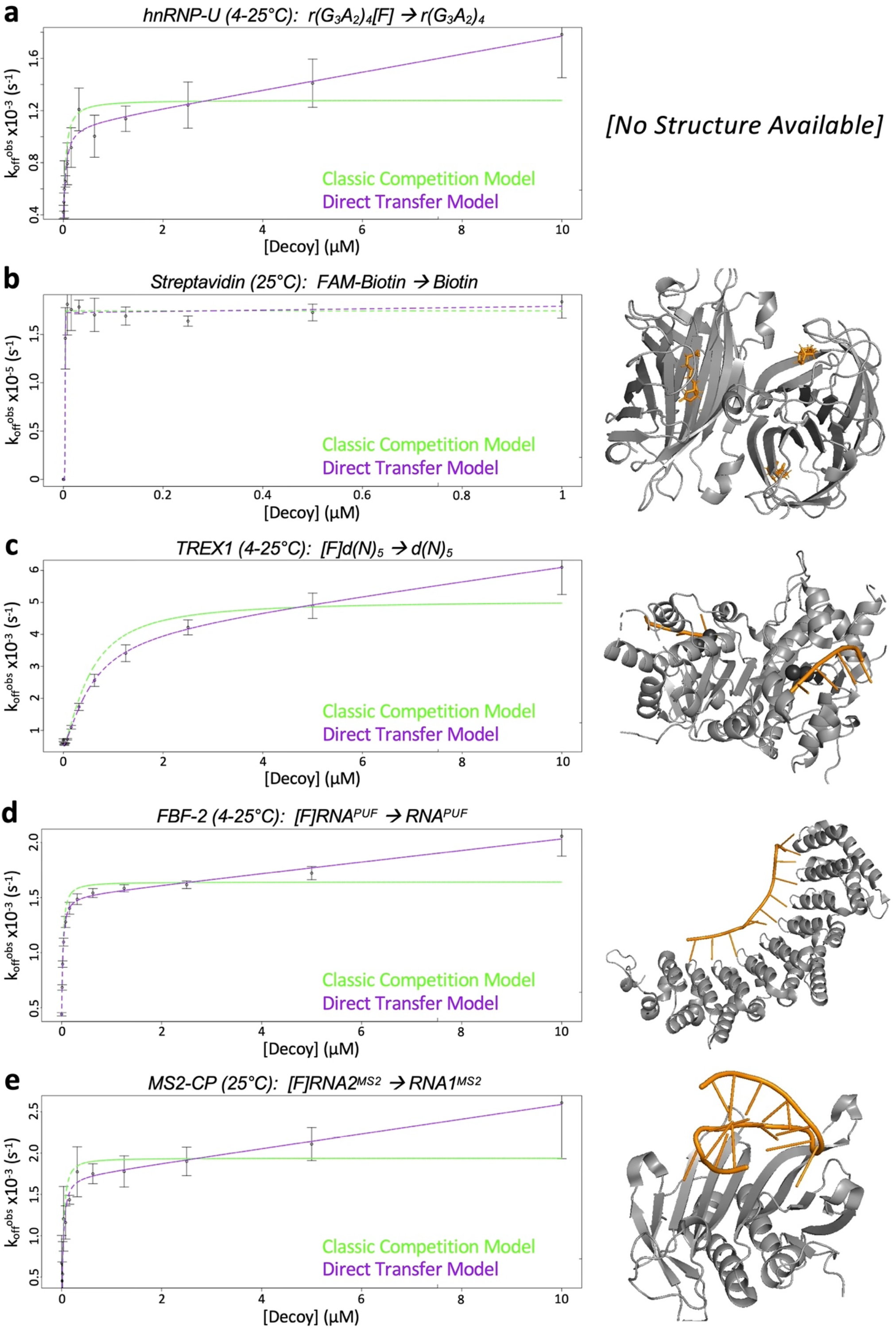
All Tested Nucleic Acid Binding Proteins Exhibit Kinetics Consistent with Direct Transfer. For several protein-polynucleotide interactions, fluorescence polarization-based competition experiments were performed and analyzed as described (Supp. Fig. 1), and the final plots of apparent prey dissociation rate (k_off_^obs^) versus decoy concentration are shown alongside crystal structures of their interactions. Plots show best-fit regression of the data with equations describing classic competition (green lines) versus direct transfer (purple lines). Graphs are from representative experiments, and error bars are mean ± SD across four technical replicates. Crystal structures show proteins as grey cartoons and nucleic acid ligands as orange cartoons/sticks. Structures in panels b-e are PDB 2IZJ, 2OA8, 3V74, and 2C51, respectively. The corresponding regression values can be found in Table 1.

Since it was unexpected that these diverse protein-nucleic acid interactions would all exhibit direct transfer, we considered possible artifactual explanations for the data. First, the assumptions of our regression and analysis approach (see Theoretical Background and Methods) include pseudo-exponential dissociation curves for reactions, so non-exponential dissociation of our complexes might produce k_off_^obs^ calculation errors that mimic direct transfer kinetics. However, our raw data clearly exhibited the expected exponential dissociation (Supp. Fig. 2). Next, some of our initial experiments (Fig. 1) were incubated at 4°C prior to reaction initiation (decoy addition) to slow apparent off-rates and reduce data variance, and these reactions warmed over time. To interrogate the effects of this temperature variation, we repeated initial experiments (Fig. 1) at constant room temperature. These results indicated no qualitative differences (Supp. Fig. 3a and Table 1), and all tested protein-polynucleotide interactions still exhibited direct transfer kinetics. Third, the methodology (Supp. Fig. 1) for the experiments of Fig. 1 used a large range of decoy concentrations. Thus, it seemed possible that variations in k_off_^obs^ across decoy concentrations could result from nonspecific polynucleotide concentration-dependent factors, such as polynucleotide aggregation or perturbation of the ionic environment of the reactions. To test this, we repeated the experiments of Fig. 1 using a non-binding carrier polynucleotide to keep total decoy polynucleotide concentration constant, while varying the ratio of binding to non-binding decoy polynucleotide. Our results (Supp. Fig. 3b and Table 1) were not qualitatively different, and all protein-polynucleotide interactions still exhibited direct transfer kinetics. Collectively, our findings suggest that direct transfer is widespread among NBPs.

**Fig. 2.**
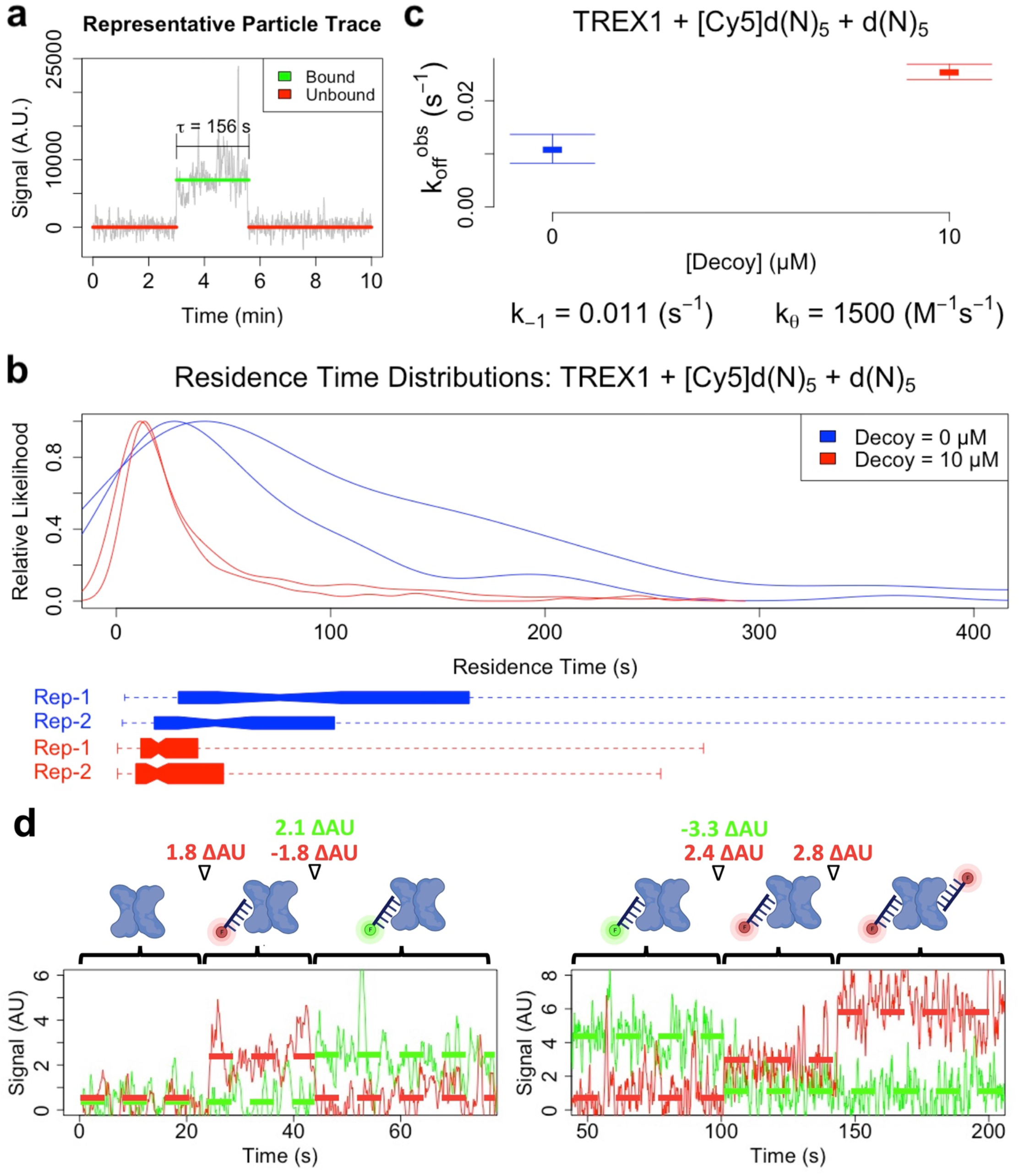
Single-Molecule Experiments Support Direct Transfer of Two DNA Molecules on TREX1. **[a-c]** Single-molecule TIRF-microscopy experiments used FLAG-tagged TREX1 conjugated to the microscope slide, Cy5-labeled prey DNA and unlabeled decoy DNA. **[a]** *Representative Particle Trace of TREX1 Binding Event*. **[b]** *Distributions of Single-Molecule Residence Times*. Slides were prepared with labeled ligand only (blue), or labeled and unlabeled ligand (red), with two replicates per condition (Rep-1 and Rep-2). Normalized kernel density plots (top graph) and box-and-whisker plots (bottom graph) of the distribution of single-particle residence times are shown for each replicate of each condition. On box-and-whisker plots, narrowed box centers indicate median, boxes define inner quartile range, and line segments define total range. **[c]** *Effect of Decoy on Apparent TREX1-ssDNA Dissociation Rate*. Apparent dissociation rates (k_off_^obs^) were calculated for each replicate/condition in panel a as the inverses of the respective distribution averages and are plotted as mean ± half-range. The dissociation rate constant (k_-1_) and direct transfer rate constant (k_θ_) were calculated from apparent dissociation rates (k_off_^obs^) as described in Methods. **[d]** *Observation Direct Transfer of Ligands on TREX1*. Single-molecule TIRF-microscopy co-localization experiments were carried out as described, using FLAG-tagged TREX1 conjugated to the microscope slide, and a mixture of Cy5- (red) and Cy3-labeled (green) ligands. Slides were prepared with trace concentrations of protein and high concentrations of both ligands, then respective binding states simultaneously monitored via dual red and green channel excitation and imaging. Example traces of ligand transfer events are shown (from n = 36). Cartoons illustrate respective binding states for TREX1 homodimer (blue), Cy3-labeled ligand (green), and Cy5-labeled ligand (red). Solid lines indicate raw signal, dashed horizontal lines indicate average signal per state, and corresponding changes in average signal between states (ΔAU) are shown at top between corresponding cartoons.

**Fig. 3.**
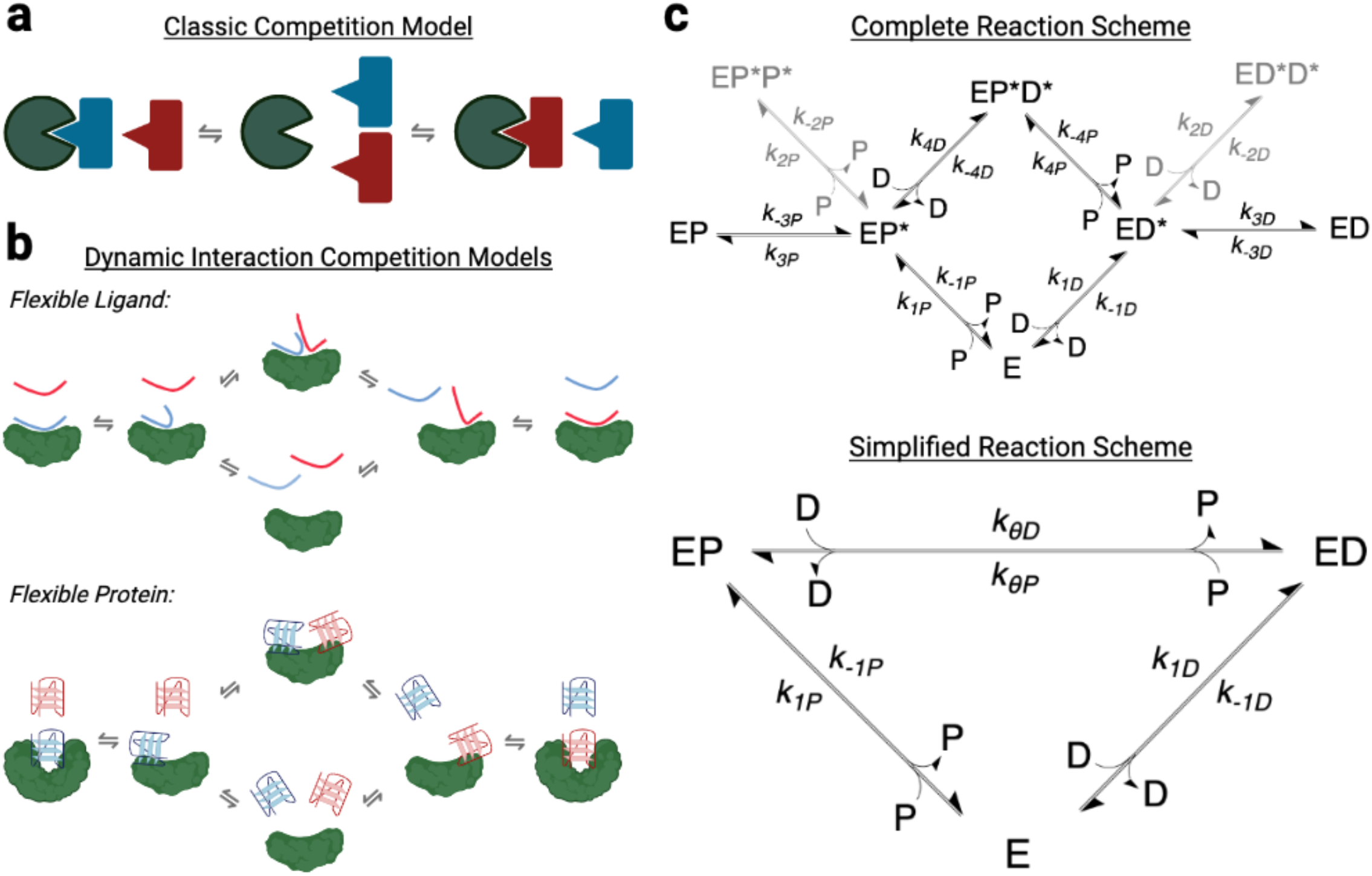
Models and Reaction Schemes for Protein-Ligand Binding Competition. **[a]** *Classic Model of Protein-Ligand Binding Competition*. The initial complex fully dissociates at a rate independent of free ligand, then the resulting free protein (green shape) can be bound opportunistically by a competing ligand (red shape), which prevents re-binding of the original ligand (blue shape). **[b]** *Dynamic Models of Protein-Polynucleotide Binding Competition*. Many nucleic acid binding reactions exhibit flexibility in associating ligands (‘Flexible Ligand’) and/or proteins (‘Flexible Protein’), as well as extensive protein-ligand contacts, so they might be better described by a dynamic model that allows the formation of short-lived ternary complexes and partial ligand interactions. Such allowances would permit competing ligands (red shapes) to influence how quickly protein (green shape) dissociates from an initially bound ligand (blue shapes), and it would also allow direct protein transfer between ligands. We term these “direct transfer” models. **[c]** *Reaction Schemes for Direct Transfer*. Complete Reaction Scheme includes protein (E), decoy (D), prey (P), and numerous rate constants (k), where conjugations of reactants are complexes, and asterisked complex components are partially associated with protein (see Eq. 1). If partially associated complexes are presumed to be highly transient, then the Simplified Reaction Scheme can describe the concentrations of remaining reactants and stable complexes (see Eq. 2).

### Single-Molecule Experiments Support Direct Transfer for TREX1

To further investigate the direct transfer model, we employed total internal reflection fluorescence (TIRF) microscopy-based single-molecule biophysics (SMB) experiments (33) as an orthogonal approach to rule out FP-dependent artifacts. We chose the TREX1 + 5-mer ssDNA interaction for these experiments because it was well behaved. Although TREX1 is a dimer, the two DNA-binding sites cannot be spanned by DNA molecules of the length tested, and it doesn’t exhibit cooperative binding for short ssDNA ligands (16, 32). Briefly, TREX1 was conjugated to a microscope slide, fluorophore-labeled oligo was flowed onto the slide with/without an excess of unlabeled oligo, and millisecond-interval movies were recorded to track individual particle binding events (Fig. 2a). Distributions of residence times across all individual binding events (Fig. 2b) were used to calculate k_off_^obs^ for each movie (Fig. 2c), and then average k_off_^obs^ values under each reaction condition were used to approximate k_-1P_ and k_θD_ (Fig. 2c). The SMB results (k_-1P_ ≈ 1.1×10^−2^ s^-1^, k_θD_ ≈ 1500 M^-1^s^-1^) were remarkably consistent with our corresponding FP data (k_-1P_ ≈ 0.7×10^−2^ s^-1^, k_θD_ ≈ 1500 M^-1^s^-1^). These findings demonstrate that direct transfer kinetics for TREX1 are not an artifact of FP-based methodology.

To directly observe transfer events, we modified the initial SMB experiments to simultaneously track the TREX1-binding states of 5-mer ssDNAs labeled with two different fluorophores. From these experiments, we identified multiple (n = 36; p < 0.01 by Monte-Carlo simulation) instances where the binding of one ssDNA coincided with the release of a differentially labeled ssDNA (Fig. 2d). We observed no instances in these experiments of more than two ligand molecules stably binding a TREX1 homodimer. We also repeated these dual-label experiments under conditions to detect Förster/fluorescence resonance energy transfer (FRET), but we were unable to detect any stable FRET signals (raw data available; see Methods). Collectively, these findings are inconsistent with any TREX1 active-sites stably binding multiple ligands, and they suggest that TREX1 can be directly transferred between ligands via an extremely short-lived ternary intermediate.

### Non-multimeric Interactions Can Exhibit Direct Transfer Kinetics

Existing reports of comparable polynucleotide transfer kinetics have concerned multimeric complexes with higher protomer-polynucleotide binding ratios, where ternary intermediates can be achieved by separate protomers binding separate ligands (1–3, 34). However, existing crystal structures of TREX1 (35), FBF-2 (19), and MS2-CP (36) bound to ligand reveal only one active site contributes to binding of each ligand, suggesting that their direct transfer must be occurring within a single protomer binding site. To confirm this implication, we performed FP-based binding curve experiments under stoichiometric conditions to determine the number of ligand molecules bound to each functional unit of our proteins. Our results for all protein-polynucleotide interactions (Supp. Fig. 4) matched the stoichiometry expected from their existing crystal structures. These findings indicate that direct transfer can occur for protein-polynucleotide interactions within a single protomer binding site.

**Fig. 4.**
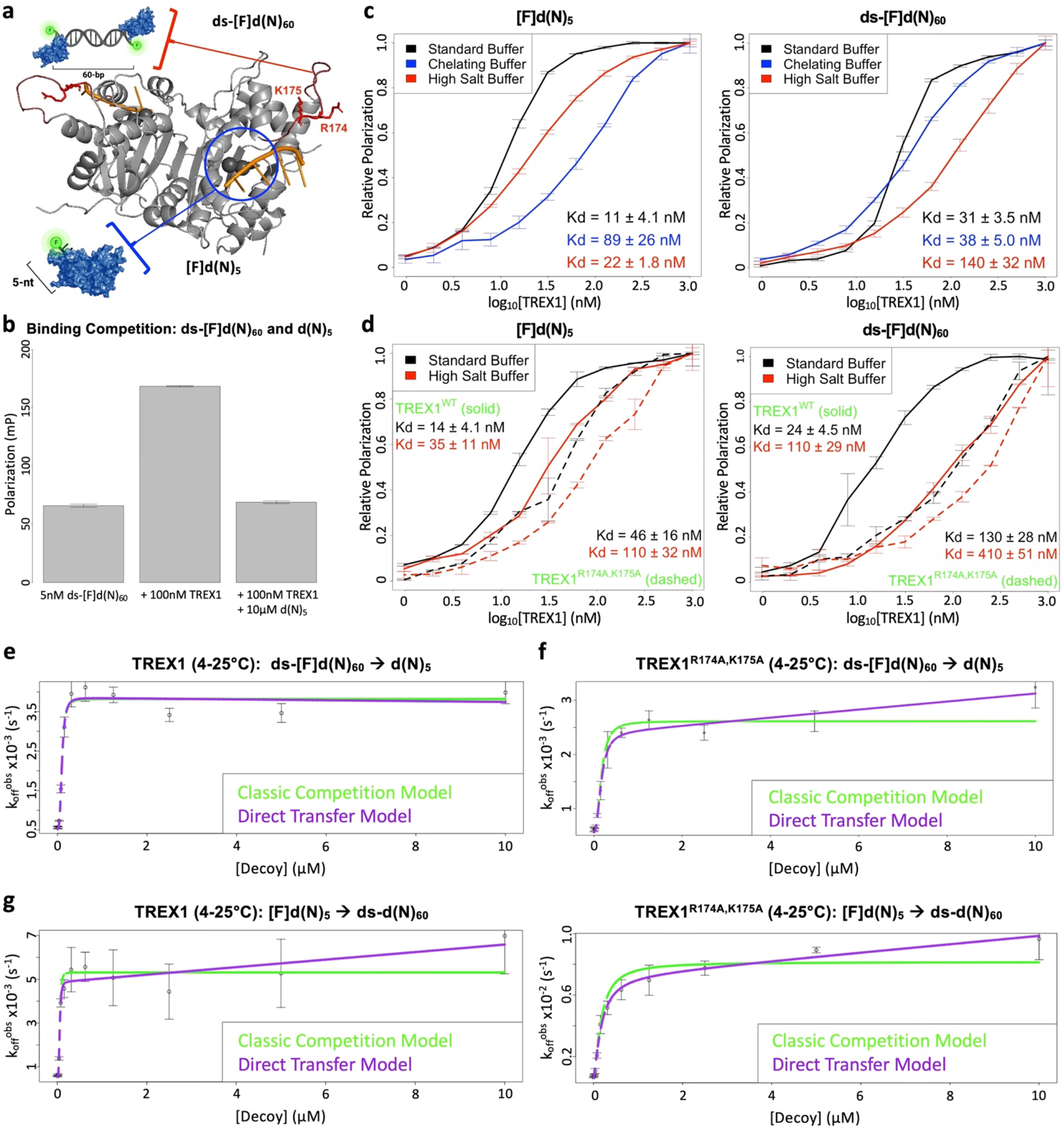
Two Competing TREX1 Ligands Exhibit Different Magnitudes of Direct Transfer. **[a]** *TREX1 Binds Long dsDNA with Extra Protein-Polynucleotide Contacts*. TREX1 (grey cartoon; ref. (16)) binds exposed 3’-hydroxyls of DNA (orange cartoon) in its nucleotide binding pocket (blue circle) with the help of divalent metal ions (black spheres). TREX1 makes additional contacts with long dsDNA through its flexible binding loop (dark red cartoon), particularly residues R174 and K175 (red sticks, labeled). We designed a 5-nt ssDNA that exclusively binds TREX1’s nucleotide binding pocket ([F]d(N)_5_), and a 60-bp dsDNA having significant additional interactions with TREX1’s flexible binding loop (ds-[F]d(N)_60_). (16). **[b]** *Two TREX1 Ligands Exhibit Competitive Binding*. Fluorescence polarization was measured for labeled 60-bp dsDNA alone (5 nM ds-[F]d(N)_60_), TREX1 with labeled 60-bp dsDNA (+ 100 nM TREX1), and TREX1 with labeled 60-bp dsDNA and unlabeled 5-nt ssDNA (+ 100 nM TREX1 + 10 µM d(N)_5_). Plot bars indicate mean ± SD from 8 technical replicates. **[c]** *Effects of Buffer Conditions on K*_*d*_^*app*^ *for Two Ligands Binding to TREX1*. Graphs are composites of three experiments with at least two technical replicates each, where error bars indicate mean ± SD, and K_d_ values are mean ± SD across the replicate experiments. **[d]** *Effects of Flexible Binding Loop Mutations on K*_*d*_^*app*^ *for Each Ligand*. Standard binding experiments with TREX1 (solid lines) and TREX1^R174A,K17A^ (dashed lines) were performed for the indicated ligands and binding buffers. Number of replicates and error bars as in [c]. **[e-g]** *Direct Transfer for the TREX1 ds[F]d(N)*_*60*_ *and d(N)*_*5*_ *Ligands Is Tunable*. FPC experiments (Supp. Fig. 1) were performed for the indicated substrates, and the data fit with equations for classic competition (green line) and direct transfer (purple line). Graphs are from representative experiments (n ≥ 3 for panels e-f, n = 1 for panel g), where error bars are mean ± SD across four technical replicates.

### The Direct Transfer Mechanism Is Revealed in TREX1 Ligand Competition

Direct transfer kinetics were ubiquitous among our tested NBPs (Fig. 1), and its occurrence with virtually identical prey and decoy ligands implicated shared protein contacts in the direct transfer mechanism. Consequently, we suspected that NBPs might depart from the classic model of ligand competition (Fig. 3a) due to intrinsically dynamic protein-polynucleotide binding interfaces that facilitate direct transfer (Fig. 3b). Indeed, as we outline in Theoretical Background, such behavior is expected to recapitulate the kinetics we observe experimentally (Fig. 3c and Supp. Fig. 1). This model for the direct transfer mechanism (Fig. 3b) suggests that its kinetics are likely influenced by the proportion of shared versus unshared protein contacts for competing ligands. TREX1 has well-characterized properties that make it suitable for mechanistically interrogating these implications. Existing crystal structures demonstrate that the 3’-termini of short ssDNA (35) and lengthier dsDNA (37) are positioned identically in TREX1’s nucleotide binding pocket (Fig. 4a), and biochemical studies indicate that lengthy dsDNA (>10 bp) has significant additional interactions with TREX1’s flexible binding loop (Fig. 4a) that don’t occur for short ssDNA (16, 38). Thus, we designed two different ligands for TREX1 (Fig. 4a): a ssDNA 5-mer, [F]d(N)_5_, which should only bind in TREX1’s nucleotide binding pocket, and a 60-bp dsDNA, ds-[F]d(N)_60_, which should bind in TREX1’s nucleotide binding pocket with an additional “foothold” on its flexible binding loop. We anticipated that the dsDNA ligand’s foothold on TREX1’s flexible binding loop would prevent efficient direct transfer of the dsDNA ligand to the ssDNA ligand, while direct transfer of ssDNA to dsDNA should occur.

We confirmed competitive binding of the two DNAs to TREX1 via FP (Fig. 4b). Next, we tested the proposed binding interactions for the two ligands. Protein-ligand interactions in the TREX1 nucleotide binding pocket occur primarily through its two divalent metal ions and hydrogen bonding with 3’-terminal nucleotides (35), while interactions with its flexible binding loop are reportedly via ionic bonding with the DNA backbone (16). We measured TREX1 binding to both ligands by FP. TREX1 binding affinity for the ssDNA ligand was significantly reduced by divalent metal ion chelation and minimally reduced by higher salt concentrations, while its affinity for the dsDNA ligand exhibited the opposite trend (Fig. 4c), supporting the binding interactions proposed for these two ligands. To further validate the binding interactions, we introduced the R174A/K175A suppression-of-function mutations (38) (Fig. 4a) to TREX1’s flexible binding loop and repeated these binding experiments. Our results (Fig. 4d) indicated that the mutant has affinity for the ssDNA ligand comparable to that of the WT protein with no change in salt sensitivity, but the mutant has significantly reduced affinity for the dsDNA ligand with ablated salt sensitivity. Collectively, these findings confirm that our two TREX1 ligands are competitive binders, that the ssDNA ligand primarily interacts with TREX1’s nucleotide binding pocket, and that the dsDNA ligand has a significant additional foothold on TREX1’s flexible binding loop.

To test if the dsDNA ligand was resistant to displacement by the ssDNA ligand, we performed FPCD experiments (Supp. Fig. 1) with dsDNA prey and ssDNA decoy (dsDNA→ssDNA). Our results (Fig. 4e and Table 1) demonstrate a decoy concentration-independent dissociation rate at high decoy concentrations. This is consistent with classic competition, indicating that direct transfer was mitigated relative to the ssDNA→ssDNA FPCD experiments (Fig. 1c), and providing our first direct evidence that protein-polynucleotide interactions can exist without detectable direct transfer. Repeating these experiments with the TREX1 flexible binding loop mutant, which should perturb the dsDNA ligand’s foothold, restored direct transfer kinetics (Fig. 4f and Table 1), consistent with our proposed mechanism that the flexible loop acts as an additional foothold for dsDNA to prevent its displacement. Finally, we expected that repeating both experiments with ssDNA ligand as prey and dsDNA ligand as decoy would follow direct transfer kinetics, since the ssDNA ligand has no unique foothold to mitigate direct transfer by the dsDNA ligand. Indeed, our results (Fig. 4g) were consistent with direct transfer kinetics for both the wild-type and mutant TREX1 enzymes. These findings support our proposed direct transfer mechanism (Fig. 3b).

### Fast Intrinsic Dissociation Is Correlated with Rapid Displacement

Our model for the direct transfer mechanism (Fig. 3b) suggests (see Theoretical Background) that direct transfer (k_θ_) and dissociation (k_-1_) rate constants should be correlated (Eq. 6.3). To test this implication, we compiled rate constant data from these studies (Table 1), concurrent studies (companion manuscript), and applicable published studies (1–3), and plotted k_θD_ as a function of k_-1P_ (Fig. 5). Our results indicate a striking correlation between the rate constants (R^2^ = 0.91), and this correlation was well maintained for only this study (R^2^ = 0.99) and only published studies (R^2^ = 0.89). Notably, outliers above the trendline in Fig. 5 involve prey ligands with presumably extensive protein contacts (G) and/or potential decoy-specific footholds (F), while interactions below the trendline tend to involve sequence-specific binding (B), potential prey-specific footholds (H), rigid prey and/or decoy ligands (C-E), and/or limited binding interactions (A & D). Taken together, these findings support the direct transfer mechanism (Fig. 3b) and its consistency among numerous independent studies.

**Fig. 5.**
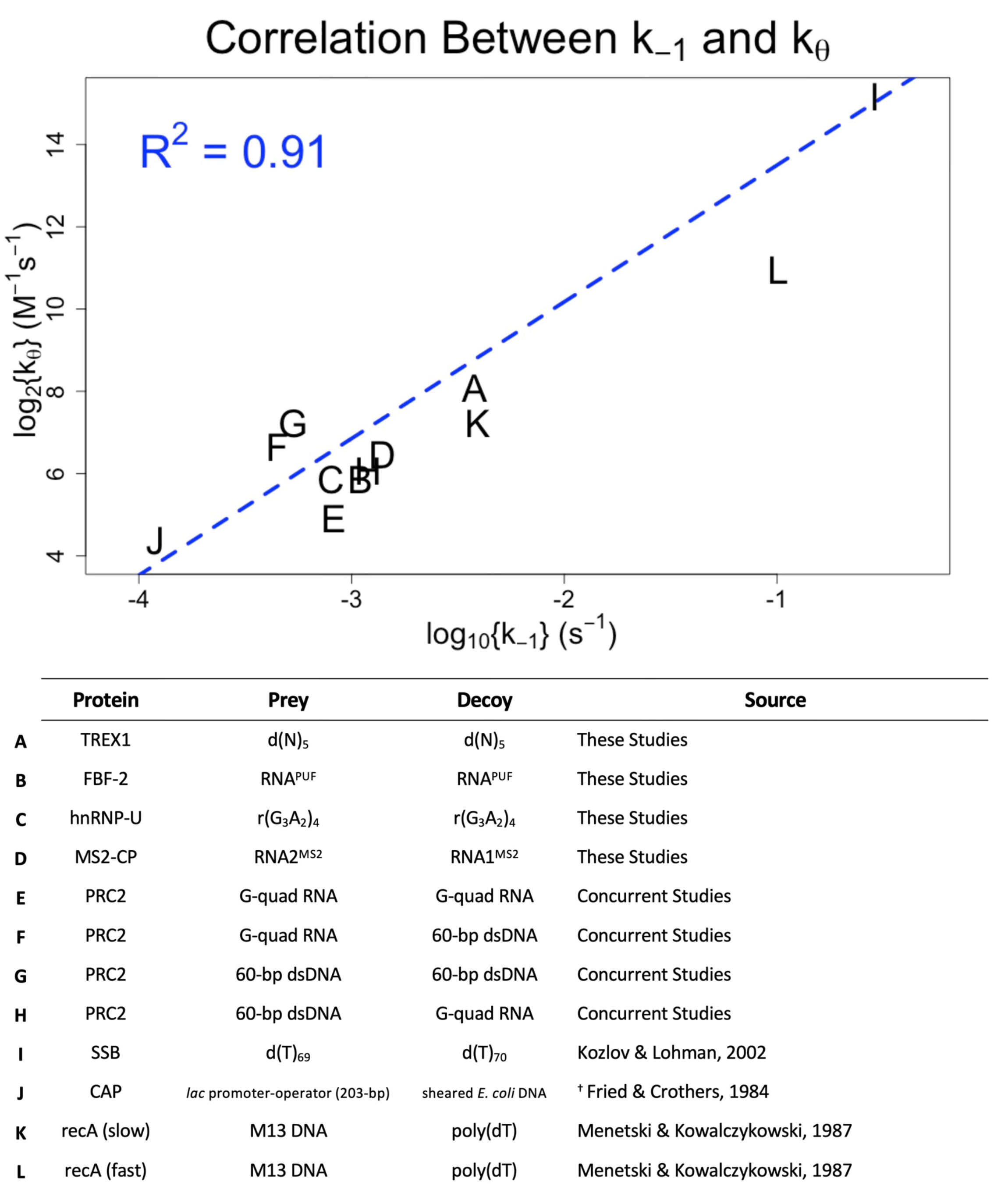
Direct Transfer Kinetics Correlated with Intrinsic Dissociation Kinetics. Direct transfer rate constants (k_θD_) are plotted as a function of intrinsic dissociation rate constants (k_-1P_) for the protein-polynucleotide interactions listed with corresponding letters below. Regression of the data (blue dashed line) was performed on linear axes with a linear 0-intercept model. Data table format mimics Table 1. Data sourced from these and concurrent studies refer to the lowest temperature competition experiments with no carrier polynucleotide, and other data are sourced as described in Methods. ^†^ Rate constant values were not explicitly stated in source; values were calculated from other source data as described in Methods.

## Discussion

Our surveyed NBPs included RNA- and DNA-binders, single- and double-stranded polynucleotide ligands, disordered and folded polynucleotide structures, variable protein-polynucleotide contact surface areas, and sequence-independent versus sequence-specific binders. Notably, while the efficiency of direct transfer varied among these interactions (Fig. 5 and Table 1), it occurred for virtually every interaction we studied. This implies that direct transfer may be a common phenomenon utilized by the majority of NBPs, though definitive conclusions will require further experimental testing. Importantly, we note that direct transfer appears to occur even between identical polynucleotides (i.e., self-competition), so NBPs with a single preferred ligand are not exempt from the consequences of direct transfer.

### Requirements for Direct Transfer

Direct transfer has been extensively studied for oligomeric nucleic acid binding proteins (1–5, 39, 40). For such proteins, it is intuitive how an incoming polynucleotide could gain a “foothold” on one of several unbound protomers to facilitate its displacement of a second polynucleotide pre-bound to a different protomer. Our findings (Fig. 1 and Supp. Fig. 4) expand the relevance of direct transfer, suggesting that it is common among RNA- and DNA-binding proteins that interact with a polynucleotide through only one binding site. Our working hypothesis is that direct transfer stems from dynamic protein-ligand contacts in a protein binding site, as shown in Fig. 3b. This implies that direct transfer fundamentally relies on (1) shared protein-ligand contacts between competitors, (2) the capacity for transient partial protein-ligand interactions, and (3) the capacity to form transient ternary complexes. In the case of TREX1, our single-molecule experiments showed multiple examples of near-simultaneous binding of one ssDNA and release of another (Fig. 2d), without stable FRET signaling, consistent with a short-lived ternary complex. Furthermore, the magnitude of direct transfer (as reflected in k_θ_) should be directly related to the extent of protein-ligand contacts, the proportion of protein contacts shared by two ligands, and the dynamicity of the protein-ligand interface(s), and it should be inversely related to the strength of contributing protein-ligand interactions. The last of these proposals is supported by the correlation between the k_-1_ and k_θ_ rate constants in Fig. 5.

The degree to which each of these properties contributes to direct transfer is an open question. Since streptavidin-biotin epitomizes extremes of these properties and does not exhibit direct transfer (Fig. 1b), we expect that there are thresholds for each of these properties, past which direct transfer cannot meaningfully occur. However, we tested high-affinity interactions with RNA hairpins (Fig. 1e) and 3-4 nt of DNA (Fig. 1c), which have very limited protein binding interfaces, and these still exhibited direct transfer, although their direct transfer efficiency (relative k_θ_/k_-1_) was lower. Finally, while direct transfer was absent for streptavidin-biotin, it may still occur for other small-molecule binders (28). Our findings suggest requisite properties for direct transfer and provide an initial framework for their quantitative boundaries, but they compel further studies with other protein-ligand interactions to better define the biophysical requirements of direct transfer.

### Considerations for Methodologies to Assess Dissociation Rate Constants and/or Mean Lifetimes

Recent studies (30, 34, 41, 42) have interrogated the related phenomenon of “facilitated dissociation” (FD), where proteins compete one-another off gene targets through a mechanism similar to how ligands compete one-another off proteins in direct transfer. Given the kinetic similarities, some implications of FD studies are echoed for direct transfer. Notably, methodologies that infer dissociation rates through “chase” of pre-formed complexes with an excess of competitor, such as the FP-based experiments used here (43, 44), should consider that the competitor may not be a neutral participant in the reaction. Making measurements across competitor concentrations may confirm classic competition, but if direct transfer is instead revealed, then extrapolation is required to derive the intrinsic dissociation rate (Supp. Fig. 1). Alternatively, methods that don’t rely on competitors to prevent reassociation could be considered, such as surface plasmon resonance (SPR) (45). SPR removes dissociated ligand from the reaction via constant buffer flow across a binding surface, which should limit direct transfer. Like “chase” experiments, single-molecule approaches that leave dissociated ligand in the reaction can have their measurements of mean complex lifetime skewed by direct transfer events causing premature complex dissociation, though the effects should be negligible under sufficiently low ligand concentrations (∼1% error with 100 nM ligand, given k_θ_/k_-1_ ≈ 10^5^ M^-1^ for most interactions). Approaches to determine the equilibrium dissociation constant (K_d_) are unlikely to have their measurements affected by direct transfer, since ligands that prematurely dissociate during direct transfer events are likely replaced by another ligand, keeping the total concentration of complex constant. This has been discussed in greater depth for the FD phenomenon (30), and the conclusions are generally applicable to direct transfer.

### Biological Relevance of Direct Transfer

Since the potential for direct transfer between polynucleotides is an intrinsic property of many nucleic acid-binding proteins, we speculate that it is likely to occur under some circumstances *in vivo*. However, it is challenging to predict which instances of direct transfer will be biologically relevant. Just because direct transfer can be measured *in vitro* doesn’t mean that it represents a mechanism integral to the protein’s function *in vivo*. Regardless, the following implications are worth consideration. First, the mean lifetime of an NBP-ligand complex may not be a fixed value, determined by the dissociation rate constant, but may instead be modulated by the concentration of other binders in its immediate environment. For example, in a nuclear domain with a high concentration of nascent transcripts, an RNA-protein interaction would be predicted to dissociate more rapidly. As another example, the high concentrations of binding sites in biological condensates or liquid-liquid phase-separated granules could promote substantial migration of nucleic acid-binding proteins within the condensate. Second, ternary complexes likely occur to some degree during polynucleotide binding competition, even if they are too unstable to be detected in equilibrium measurements. So, future hypotheses relying on ternary complex formation should not be discounted based solely on a lack of experimentally detectable ternary complexes for an NBP. Third, for large NBP ligands (e.g., chromatin), translocation of the NBP between two sites on the ligand molecule need not be mediated by protein sliding or free diffusion, but it might instead occur by intramolecular direct transfer when sites become spatially proximal (e.g., due to folded/looped chromatin architecture). Finally, one polynucleotide species can still facilitate binding of another polynucleotide species to their NBP partner (and the NBP’s activity on that ligand), even if they actively compete for binding to that NBP. In fact, in concurrent studies of PRC2 (companion manuscript) we provide quantitative evidence that such recruitment can occur if certain conditions in the reaction environment are met, and that this can allow tunable RNA-mediated regulation of chromatin binders.

In terms of both efficiency and magnitude, TREX1 exhibits average direct transfer (Fig. 5), but there is no readily apparent role for direct transfer in promoting TREX1’s degradation of extranuclear DNA to prevent nucleic acid sensing (10). However, TREX1 is purportedly capable of “sliding” along DNA for efficient substrate binding and exonucleolytic degradation (16), and direct transfer could be an alternative or additional way to reduce the dimensionality of an otherwise 3D search process (27, 28). At minimum, the observation that TREX1’s flexible loop can drastically modulate its direct transfer kinetics (Fig. 4e-f) demonstrates the flexible loop’s critical contribution to TREX1 dsDNA binding affinity, consistent with recent studies (16). In contrast to TREX1, PRC2 exists as a clear trendline outlier (Fig. 5), and our concurrent studies (companion manuscript) suggest that direct transfer may explain some of PRC2’s biological function: RNA-mediated regulation of PRC2’s histone methyltransferase activity on nucleosomes. Thus, direct transfer likely serves a biological role for PRC2. Indeed, we suspect RNA-binding chromatin-associated proteins may generally exploit direct transfer to target their genetic loci (see companion manuscript).

Finally, what *in vivo* approaches might be used to test the biological role of a protein’s direct transfer? Ideally, one would specifically ablate the capacity for a protein to perform direct transfer between two ligands without affecting their independent binding, presumably though mutation of the protein or polynucleotide sequences, and then compare phenotypes. However, since direct transfer appears to be mediated by protein contacts shared between ligand species (Fig. 3b), modulating direct transfer might necessarily perturb the ligands’ independent binding. In that case, the mutant phenotype could not be specifically attributed to direct transfer versus affinity perturbations. Similarly, since unique protein contacts between competing ligands act as “footholds” that should affect direct transfer, trying to modify shared versus unshared protein contacts to achieve similar binding affinities with discrepant direct transfer kinetics could prove problematic. However, we look forward to future advances that may allow direct *in vivo* interrogation of direct transfer biology.

### Theoretical background

#### Dynamic Protein-Ligand Interactions Produce Direct Transfer Kinetics

Inter-ligand competition for dynamic protein-ligand interactions (Fig. 3b) can be generalized with a reaction scheme (Fig. 3c – Complete Reaction Scheme) that is quantitatively described by a system of differential equations (Eq. 1). If the partial interaction intermediates are presumed to be highly transient (as our data suggest), such that ([EP*],[EP*P*],[EP*D*],[ED*],[ED*D*]) << ([E],[EP],[ED],[P],[D]), then the stable reactants can be modeled with a simplified reaction scheme (Fig. 3c – Simplified Reaction Scheme) and system of equations (Eq. 2). Consider, then, a two-phase reaction under this scheme (via Eq. 2) where the pre-reaction state (1) is [E_T_] >> K_dP_, [P_T_] < [E_T_], and [D_T_] = 0 in equilibrium such that [EP] ≈ [P_T_], then reaction initiation (2) occurs by decoy addition such that [EP]_0_ ≈ [P_T_], [E]_0_ ≈ [E_T_] – [P_T_], [D]_0_ = [D_T_], and all other reactants are approximately 0. In such a reaction, if [D_T_] >> [E_T_] and [P]_0→∞_ ≈ 0 (prey liberated from protein-prey complex after initiation is lost from reaction), then the reaction obeys Eq. 3.1. Similarly, if [D_T_] >> K_dD_ + [E_T_] such that [E]_0→∞_ ≈ 0 and k_θD_ [EP]_t_ [D]_t_ + k_-1P_ [EP]_t_ >> k_θP_ [ED]_t_ [P]_t_, then the reaction is well approximated by Eq. 3.1. Solving the differential equation (Eq. 3.1) yields Eq. 3.2, which is a function for exponential dissociation. Thus, for [E_T_] ≥ K_dD_ the proposed two-phase direct transfer reactions obey Eq. 4.1 with exponential dissociation. If classic competition applies (k_θP_ = k_θD_ = 0), this reduces to Eq. 4.2.

It is evident from Eq. 2 that Eq. 4 should decrease farther below its limit as [D_T_]→0, and Eq. 5 is an adjustment to Eq. 4 that should approximate this behavior. To test this approximation’s efficacy, we simulated reactions over a range of initial decoy concentrations (via Eq. 2) using combinations of rate constant values relevant to our chosen protein-polynucleotide interactions, then used Eq. 5 to analyze the simulated reaction sets in a format mimicking our empirical experiments (see Methods). The results of our analysis (Supp. Table 1) indicate this approach provides highly accurate rate constant determinations and model classifications for the proposed reaction scheme (Fig. 3c – Simplified Reaction Scheme), which are applicable to our experimental strategy (Supp. Fig. 1).

#### The Proposed Mechanism for Direct Transfer Predicts Correlated Rate Constants

Though not explicitly indicated by our simplified reaction scheme (Fig. 3c) for direct transfer, it is notable from our complete reaction scheme (Fig. 3c) that the displacement of one ligand molecule by another is an opportunistic process at the sub-molecular scale. Thus, our direct transfer rate constant for decoy (k_θD_) should be well-correlated to the prey’s dissociation rate constant (k_-1P_), the decoy’s association (k_1D_) and dissociation (k_-1D_) rate constants, and the protein-polynucleotide interaction dynamicity (δ) from protein/polynucleotide flexibility and prey-decoy binding overlap. Eq. 6.1 describes this general relationship. If, for most interactions, k_1D_ varies minimally, then k_1D_ is reconciled with an arbitrary constant (a_1_) and Eq. 6.1 reduces to Eq. 6.2. If additionally, k_-1D_ < k_-1P_ such that complete prey dissociation predominately follows partial decoy association, then k_-1D_ can be ignored and Eq. 6.2 reduces to Eq. 6.3.

## MATERIALS & METHODS

### Purification of Proteins

Recombinant human hnRNP-U_673-825_ protein (RNA binding domain) with an N-terminal fusion of His-tagged Maltose Binding Protein (HisMBP) was purified by Otto Kletzien (University of Colorado Boulder, Department of Biochemistry, Batey Lab) as previously detailed (8). Protomer concentration was determined by spectroscopy with ε_280_ = 99,240 M^-1^cm^-1^.

Streptavidin was purchased (Sigma Aldrich #S0677-5MG, Lot #SLCJ1102) as dry powder, resuspended in buffer, and its active-site (protomer) concentration determined by spectroscopy with ε_280_ = 40,300 M^-1^cm^-1^.

Recombinant murine TREX1_1-242_ protein (catalytic core) was overexpressed and purified as previously detailed (46). Briefly, protomers were expressed as a fusion protein using a pLM303x vector that encodes maltose-binding protein (MBP) linked N-terminally to TREX1 with a rhinovirus 3C protease (PreScission Protease) recognition site. Plasmid was transformed into *E. coli* Rosetta2(DE3) cells (Novagen) for overexpression, cells lysed with an Emulsiflex C3 homogenizer (Avestin), then protein purified via sequential amylose column chromatography, overnight protease cleavage, and phosphocellulose (p-cell) column chromatography. The reported wild-type plasmid was subjected to site-directed mutagenesis to introduce R174A/K175A mutations or a C-terminal FLAG tag (DYKDDDDK), then the plasmids’ identities were validated by sequencing. The mutated plasmids were used to obtain mutant and FLAG-tagged protein similarly to wild-type protein, with the exception that mutant protein did not bind the p-cell column and was left as an MBP + TREX1 mixture. Preparations were determined via SDS-PAGE to be >95% purity. Active-site (protomer) concentrations were determined by spectroscopy with ε_280_ = 24,142 M^-1^cm^-1^ (TREX1 and TREX1-FLAG) or ε_280_ = 90,300 M^-1^cm^-1^ (MBP-TREX1^R174A,K175A^), and the equivalency of TREX1 concentrations between wild-type and mutant preparations was validated by SDS-PAGE.

Recombinant *C. elegans* FBF-2_164-575_ protein (RNA binding domain) was purified by Chen Qiu (National Institute of Environmental Health and Safety, Lab of Traci Hall) as previously described (19). Reported concentrations are for active-sites (monomer), unless otherwise indicated.

For recombinant MS2-CP^V75E,A81G^ protein (mutant without capsid assembly (25)), a pMAL-c6T vector (NEB #N0378S) was altered by site-directed mutagenesis to replace the TEV protease recognition site with a rhinovirus 3C protease (PreScission Protease) recognition site. This produced the “pMALcPP” vector, which encodes HisMBP-protein fusions. Then, we used site-directed mutagenesis to introduce the V75E and A81G mutations into a yeast expression vector containing the wild-type MS2-CP gene fragment (provided by Roy Parker Lab, University of Colorado Boulder, Department of Biochemistry). The MS2-CP^V75E,A81G^ gene fragment was subcloned into our pMALcPP vector, and we confirmed by sequencing that the resulting plasmid encoded a HisMBP–MS2-CP^V75E,A81G^ fusion with rhinovirus 3C protease (PreScission Protease) linker. Then, plasmid was transformed into *E. coli* BL21(DE3) cells (NEB). The transformed cells were used to inoculate 20 mL media (LB + 100 µg/mL ampicillin) and incubated overnight at 37°C/200 rpm until A_600_ ≈ 5.0. The starter culture was diluted in 1 L fresh media so that A_600_ ≈ 0.1, incubated at 37°C/200 rpm until A_600_ ≈ 0.8 (∼2 h), induced with 0.5 mM IPTG, and incubated overnight at 16°C/200rpm. Induced cells were pelleted by centrifugation (4,000G/4°C/20 min), resuspended in 50 mL Amylose A Buffer (20 mM TRIS pH 7.5 at 25°C, 200 mM NaCl, 1 mM EDTA) + 1 Pierce Protease Inhibitor Tablet (Thermo Scientific #A32965) + 50 mg lysozyme, then lysed with an Emulsiflex C3 homogenizer (Avestin) at 15,000-18,000 psi. Lysate was clarified by centrifugation (27,000x g/4°C/30 min), then supernatant was collected. A low-pressure chromatography column connected to a peristaltic pump (5 mL/min flow rate) was prepared with 5 mL amylose resin (NEB #E8021S), followed by equilibration with 50 mL Amylose A Buffer, supernatant application, washing with 300 mL Amylose A Buffer, and elution (bulk collection) with 50 mL Amylose B Buffer (Amylose A Buffer + 10 mM maltose). To the eluent was added 1.0 mg of Prescission Protease, and the solution was loaded into 10 kDa-cutoff SnakeSkin Dialysis Tubing (Thermo Scientific #68100) and dialyzed overnight at 4°C in Nickel A Buffer (50 mM NaH_2_PO_4_ pH 8.0 at 25°C, 300 mM NaCl, 10 mM imidazole). A low-pressure chromatography column connected to a peristaltic pump (2 mL/min flow rate) was prepared with 15 mL nickel resin (Qiagen #30230), followed by equilibration with 50 mL Nickel A Buffer, application of the dialyzed sample (w/ 5 mL fraction collection), and washing with 50 mL Nickel A Buffer (w/ 5 mL fraction collection). A_280_ of each fraction was determined, and the protein-rich fractions pooled. SDS-PAGE indicated >95% purity. Active-site (homodimer) concentrations were determined by spectroscopy with ε_280_ = 34,000 M^-1^cm^-1^.

### Preparation of Ligands

All oligonucleotides were ordered from IDT (Coralville, IA), and their sequences in IDT syntax are provided (Supp. Table 2). FAM-biotin (#53606-1MG-F) and biotin (#B4501-100MG) were ordered from Sigma Aldrich. For dsDNA constructs, complementary oligos ordered from IDT were mixed at 5 µM (prey) or 300 µM (decoy) each in annealing buffer (50 mM TRIS pH 7.5 at 25°C, 200 mM NaCl) and subjected to a thermocycler program (95°C for 10 min, 95→4°C at 0.5 °C/min, hold at 4°C) for annealing, and annealing was then confirmed via Native-PAGE. Concentrations of all ligands were confirmed spectroscopically using manufacturer-provided extinction coefficients.

### Binding Buffer Compositions

BB1 is 50 mM TRIS (pH 7.5 at 25°C), 25 mM KCl, 2.5 mM MgCl_2_, 0.1 mM ZnCl_2_, 0.1 mg/mL BSA, 5% v/v glycerol, 2 mM 2-mercaptoethanol. Proteins = hnRNP-U and streptavidin.

BB2 (TREX1 ‘Standard Buffer’) is 20 mM TRIS (pH 7.5 at 25°C), 5 mM CaCl_2_, 2 mM DTT, 0.1 mg/mL BSA. Protein = TREX1.

BB3 (TREX1 ‘High Salt Buffer’) is 20 mM TRIS (pH 7.5 at 25°C), 5 mM CaCl_2_, 200 mM NaCl, 2 mM DTT, 0.1 mg/mL BSA. Protein = TREX1.

BB4 (TREX1 ‘Chelating Buffer’) is 20 mM TRIS (pH 7.5 at 25°C), 5 mM CaCl_2_, 5 mM EDTA, 2 mM DTT, 0.1 mg/mL BSA. Protein = TREX1.

BB5 is 10 mM HEPES (pH 7.5 at 25°C), 50 mM NaCl, 0.01% v/v Tween-20, 2 mM DTT, 0.1 mg/mL BSA. Protein = FBF-2.

BB6 is 100 mM TRIS (pH 7.5 at 25°C), 10 mM MgCl_2_, 80 mM KCl, 0.1 mg/mL BSA. Protein = MS2-CP.

### FP-Based K_d_ Determination

Pre-reaction mix was prepared with 5 nM prey molecule in the indicated binding buffer (see Binding Buffer Compositions), then dispensed in 36 µL volumes into the wells of a 384-well black microplate (Corning #3575). Protein was prepared at 10X the reported concentrations via serial dilution in binding buffer. Binding reactions were initiated by addition of 4 µL of the respective protein concentration to the corresponding pre-reaction mix and then incubated for 30 min at room temperature. Wells with binding buffer only were also included for blanking. Fluorescence polarization readings were then taken for 30 min in 30 s intervals with a TECAN Spark microplate reader (Ex = 481 ± 20 nm, Em = 526 ± 20 nm). Each experiment had 2 or 4 technical replicates per protein concentration (as indicated), and at least three experiments were performed per protein-polynucleotide interaction. Protein concentrations are defined previously (see Purification of Proteins).

Raw data were analyzed in R v4.1.1 with the FPalyze function (FPalyze v1.3.0 package). Briefly, polarization versus time data were calculated for each reaction, the last 10 data points for each reaction were averaged to generate an equilibrium polarization value, and equilibrium polarization values were plotted as a function of protein concentration. Plot data were regressed with Eq. 7 to calculate K_dapp_ for the interaction.

### FP-Based Competitive Dissociation Experiments

Pre-reaction mix was prepared with 5 nM prey molecule and protein ≥ 2x K_dPapp_ (at 25°C) in the indicated binding buffer (see Binding Buffer Compositions), then dispensed in 36 µL volumes into the wells of a 384-well black microplate (Corning #3575). Decoy was prepared at 10X the reported concentrations via serial dilution in binding buffer or the respective carrier polynucleotide (Table 1) at a concentration equal to the highest decoy concentration. Pre-reaction mix and decoy dilutions were then incubated at the indicated temperature until thermal and binding equilibrium (4°C/90 min, 25°C/30 min, or 37°C/30 min). Competitive dissociation reactions were initiated by addition of 4 µL of the respective decoy concentration to the corresponding pre-reaction mix, then fluorescence polarization readings were immediately taken (the delay between initiation of the first reactions and the first polarization reading was ∼90 s) at the indicated temperature (25°C or 37°C) for 120 min (the streptavidin-biotin dissociation rate was so slow that the reactions had to be extended to 24 h with the plate placed in a humidity cassette to mitigate evaporation) in 30 s intervals with a TECAN Spark microplate reader (E_x_ = 481 ± 20 nm, E_m_ = 526 ± 20 nm). Each experiment had 4 technical replicates per decoy concentration, and at least three experiments were performed per protein-polynucleotide interaction unless otherwise indicated. Specific protein concentrations used were (Protein–Prey = [Protein]): TREX1^R174A,K175A^–ds-[F]d(N)_60_ = 250 nM, All Others = 100 nM. Protein concentrations were determined as described (see Purification of Proteins).

Raw data was analyzed in R v4.1.1 with the FPalyze function (FPalyze v1.3.0 package). Briefly, polarization versus time data were calculated for each reaction, the polarization data were normalized to the maximum and minimum polarization across all reactions, and each normalized reaction was fit with an exponential dissociation function (Eq. 8.1) to determine k_off_^obs^ (Eq. 8.2). Then k_off_^obs^ values were plotted as a function of decoy concentration. Plot data (with background k_off_^obs^ subtracted to mitigate temperature effects on polarization) were regressed via Eq. 5.1 then Eq. 5.2 with tuning parameters constrained to the Eq. 5.1 solutions, and the regression models were compared with the Bayesian Information Criterion (BIC) (47). Rate constants (k_-1P_ and/or k_θD_) were determined from the best-performing regression model. If minimum polarization was not reached during competition experiments (e.g., due to a weak competitor), then it was manually defined with minimum polarization data from corresponding binding curve data (see FP-Based K_d_ Determination).

### FP-Based Stoichiometry Experiments

Binding curve experiments were performed as described above (see FP-Based K_d_ Determination), with a few exceptions. (1) The prey concentration was 250 nM. (2) Single experiments were performed with 4 technical replicates. (3) Protein concentrations were defined by functional units (TREX1 = homodimer, FBF-2 = monomer, MS2-CP = homodimer) instead of active-sites. (4) Plot data were regressed with Eq. 9 to determine ligand-protein stoichiometry.

### Accuracy Metrics for Rate Constant Determination Strategy

Direct transfer reactions (Eq. 2) were simulated and analyzed in R v4.1.1 with a custom script (see Software, Data, and Materials Availability). Briefly, reaction time (t), [E_T_], [P_T_], [D_T_], K_dP_, K_dD_, k_-1P_, k_-1D_, k_θP_, k_θD_, start-read delay time (t_0_), read frequency (Δt), coefficient of variation versus [EP]_0_ to use for Gaussian error (σ), number of replicate data sets to generate (N), and whether to provide the true baseline signal during analysis were user-provided. Then, initial conditions were calculated via Eq. 10, the system of differential equations (Eq. 2) was solved by numerical integration, simulated reactions were sampled (t_0_ and Δt) to mimic instrument readings, sampled reaction values were used to simulate numerous (N) erred (σ) data sets with 4 technical replicates each, each simulated data set was analyzed with the FPalyze function (FPalyze v1.3.0 package), and accuracy metrics were calculated across successfully analyzed data sets. To automate analyses, FPalyze made unsupervised estimates of Eq. 5 parameters to guide regression. Consequently, regression sometimes failed due to inadequate estimates, and data sets had to be excluded from accuracy metric calculations. By default, t = 120 min, [E_T_] = 3x K_dP_, [P_T_] = 5 nM, [D_T_] = 10x(2^0^,2^-1^,2^-2^,2^-3^,2^-4^,2^-5^,2^-6^,2^-7^,2^-8^,2^-9^,2^-10^,0) µM, t_0_ = 90 s, Δt = 30 s, σ = 0.05, N = 20, and true baseline signal was not provided to FPalyze.

### TIRF Microscopy-Based Single-Molecule Experiments

PEG-biotin coated microscope slides were prepared as previously described (33). Slides were (1) washed twice with 200 µL Millipore water (mpH2O), (2) twice with 200 µL reaction buffer (BB2 + 0.05% v/v NP-40), (3) incubated for 5 min with a mix of 0.2 mg/mL streptavidin and 0.8 mg/mL BSA in reaction buffer, (4) washed twice with 200 µL reaction buffer, (5) incubated for 5 min with 25 pM (single-label) or 5 pM (dual-label) biotin-tagged α-FLAG monoclonal antibody (ThermoFisher Scientific #MA1-91878-BTIN, Lot #XB341977) in reaction buffer, (6) washed twice with 200 µL reaction buffer, (7) incubated for 5 min with 250 nM TREX1-FLAG protein in reaction buffer, and (8) washed twice with 200 µL reaction buffer. This generated slides with TREX1 conjugated to their surface.

For single-label experiments, TREX1-conjugated slides were treated with 50 pM [Cy5]d(N)_5_ prey ± 10 µM d(N)_5_ decoy in imaging buffer (3 mM Trolox, 1% v/v glucose, 1 mg/mL glucose oxidase, and 0.1 mg/mL catalase in reaction buffer), then immediately imaged with 640 nm laser excitation and a red wavelength bandpass camera filter. Data herein are from a single experiment with two replicates per decoy concentration: no-decoy (rep-1) = 10 min movie with 300 ms exposure, no-decoy (rep-2) = 10 min movie with 150 ms exposure, high-decoy (rep-1) = 10 min movie with 300 ms exposure, high-decoy (rep-2) = 10 min movie with 150 ms exposure.

For dual-label experiments, TREX1-conjugated slides were photobleached with a 532 nm laser for 10 min, treated with 10 nM [Cy5]d(N)_5_ + 10 nM [Cy3]d(N)_5_ in imaging buffer (3 mM Trolox, 1% v/v glucose, 1 mg/mL glucose oxidase, and 0.1 mg/mL catalase in reaction buffer), then immediately imaged. Co-localization experiments used 532 and 640 nm laser excitation with green and red wavelength bandpass camera filters, and FRET experiments used 640 nm laser excitation with green and red wavelength bandpass camera filters. Co-localization data are from a single experiment with four replicate 5 min movies at 150 ms exposure. FRET data are from a single experiment with four replicate 5 min movies at 150 ms exposure.

For single-label experiments, each replicate was analyzed in R v4.1.1 with the SMBalyze 2.0.4 package. Briefly, (1) identities of the tiff stacked image files were blinded with the blind.input function, (2) the id.spots function was used to create a composite image, identify particles, and calculate signal intensity over time for each particle, and (3) the refine.particles function was used to manually validate particle traces, refine binding event selections, and export residence times for all binding events. The numbers of particles selected for residence time calculations during refinement were n = 39 (no-decoy rep-1), n = 44 (no-decoy rep-2), n = 70 (high-decoy rep-1), and n = 88 (high-decoy rep-2). The residence times were then analyzed in R v4.1.1 with a custom script to calculate k_off_^obs^ for each replicate, and to calculate k_-1P_ and k_θD_ from k_off_^obs^ data. Raw images and RData output files from each step are freely available (see Software, Data, and Materials Availability).

For dual-label experiments, each replicate was analyzed in R v4.1.1 with the SMBalyze 2.0.5 package. Briefly, (1) broad-wavelength control images were used to align red and green camera images with the FRET.align function, (2) the FRET.id function was used to create composite images, identify particles, pair particles, and calculate signal intensity over time for each particle, and (3) the FRET.refine function was used to manually validate particle traces and export data for all events. For co-localization experiment data, the number of direct transfer events identified from each replicate was n = 8 (Co-loc, r1), n = 5 (Co-loc, r2), n = 12 (Co-loc, r3), n = 11 (Co-loc, r4). Raw images and RData output files from each step are freely available (see Software, Data, and Materials Availability). For FRET experimental data, raw images are freely available (see Software, Data, and Materials Availability).

### Single-Molecule Statistical Analysis for Direct Transfer Events

Single-molecule binding states for dual-label experiments were simulated and analyzed in R v4.1.1 with a custom script (see Software, Data, and Materials Availability). Briefly, reaction time (t), exposure time (Δt), [E_T_], total concentrations of the two fluorophore labeled ligands ([L1_T_] and [L2_T_]), K_d_, k_-1_, k_θ_, number of signal traces (n), protomers in functional protein unit (p), number of data sets (m), and trace signals’ standard deviation relative to single state changes (σ) were user-provided. Equilibrium conditions were calculated via Eq. 11, and then the initial binding states for (n)x(p) particles were randomly sampled via the sample function (R base) using the equilibrium condition concentrations as weights. Binding states for each particle were iteratively updated at each time point via similar random sampling with weights defined by the probabilities of transitioning to other particle binding states within one exposure time (Δt). These probabilities were calculated via the pexp function (stats package) using Δt as the quotient parameter and the corresponding λ from Eq. 12 as the rate parameter. The probabilities of no state change for a particle during the exposure time was calculated as the residual probability. Binding states for particles were converted into arbitrary signal units for two different visualization channels, corresponding to the two different fluorophores on the ligand, and error was added to the particles’ signal traces via the rnorm function (stats package) parameterized by σ. Particles’ signal traces (in both channels) were then randomly combined into n sets of p, to simulate cumulative signal from protomers (p) in a functional protein. The upper times between every anticorrelated state change across channels were calculated for every protein molecule. This whole workflow was repeated iteratively to produce m replicate simulations of our data. Parameter settings for our script were matched to experimental conditions: t = 300 s, Δt = 150 ms, [E_T_] = 5 pM, [L1_T_] = 10 nM, [L2_T_] = 10 nM, K_d_ = 8.9 nM, k_-1_ = 1.1×10^−2^ s^-1^, k_θ_ = 0 or 1500 M^-1^s^-1^, n = 2558, m = 100, p = 2, and σ = 0.1.

Cumulative probability distributions (Supp. Fig. 5a) were generated by calculating the proportion of m simulations that had N or less anticorrelated state changes occur within various time thresholds (DTΔ). Our findings (Supp. Fig. 5 – right) indicate that we likely classified exchange events as anti-correlated state changes occurring within two (not one) exposure time frames, consistent with the signal-to-noise in our experimental data conferring error onto the time calls for state changes. Notably, 0/100 simulated experiments under a classic model (Supp. Fig. 5 – left) produced our observed number of apparent exchange events at this threshold (DTΔ ≤ 0.30 s), indicating the classic model is rejectable with p < 0.01.

### Rate Constant Correlation Analysis

Data sourced from these studies (Fig. 5 – A-D) were the average values (Table 1) from initial experiments (Fig. 2). Data sourced from concurrent studies on PRC2 (Fig. 5 – E-H) are from Figure 3 of the accompanying manuscript. The first published data (Fig. 5 – I) are taken from Table 2 of its reference (1). The second published data (Fig. 5 – J) are calculated from Table 1 values of reference (2), where k_-1P_ is the dissociation rate at the lowest DNA concentration and k_θD_ is the rate of change in dissociation rate between the highest and lowest DNA concentrations. The last published data (Fig. 5 – K-L) are taken from Figure 2 of reference (3).

Rate constants from the indicated sources were regressed with linear axes and a 0-intercept constraint in R v4.1.1 using the lm function (R base). Regression values were R^2^ = 0.91 and m = 1.2×10^5^ M^-1^ for all data (Fig. 5 – A-L), R^2^ = 0.91 and m = 1.2×10^5^ M^-1^ for these and published data only (Fig. 5 – A-D & I-L), R^2^ = 0.99 and m = 0.69×10^5^ M^-1^ for these data only (Fig. 5 – A-D), and R^2^ = 0.89 and m = 1.2×10^5^ M^-1^ for published data only (Fig. 5 – I-L).

### Diagram, Reaction Scheme, and Figure Generation

Diagrams were prepared with BioRender, reaction schemes were prepared with ChemDraw v21.0.0 (Perkin Elmer), tables were prepared with Word (Microsoft), graphs were prepared with R v4.1.1, protein structures were prepared in PyMOL v2.5.2 (Schrodinger), and figures were assembled in PowerPoint (Microsoft). The modeled murine TREX1 structure in Fig. 4 was taken from prior work (16).

### Software, Data, and Materials Availability

GitHub hosts the FPalyze (github.com/whemphil/FPalyze) and SMBalyze (github.com/whemphil/SMBalyze) R packages. The custom scripts referenced in these methods are available on GitHub (github.com/whemphil/Direct-Transfer_Manuscript). For the single-molecule experiments, raw movie files and SMBalyze output files have been uploaded to Zenodo (doi.org/10.5281/zenodo.7383172). Our pMALcPP vector and pMALcPP/MS2-CP plasmids will be deposited to Addgene.

### Equations

For Eq. 1.1-10, terms are defined in Fig. 3c – Complete Reaction Scheme, equations give rates of change for indicated reactants as a function of time (t), and bracketed terms indicate concentrations.

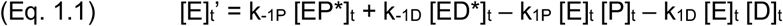

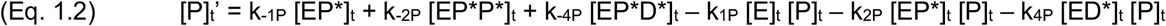

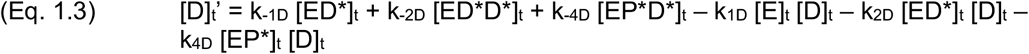

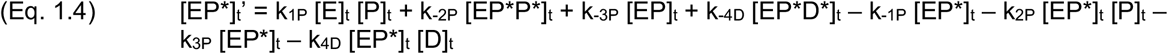

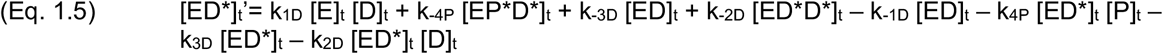

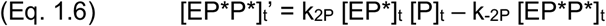

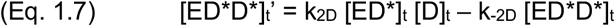

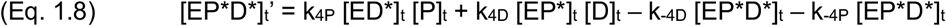

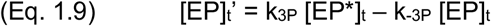

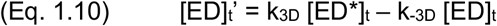

For Eq. 2.1-5, terms are defined in Fig. 3c – Simplified Reaction Scheme, equations give rates of change for indicated reactants as a function of time (t), and bracketed terms indicate concentrations. For Eq. 2.6-10, apply Eq. 2.1-5 notation.

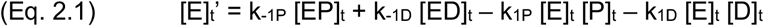

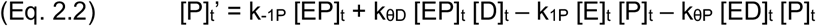

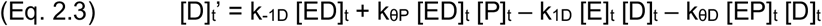

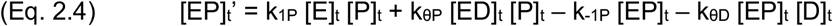

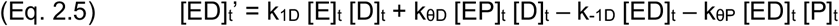

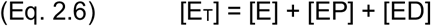

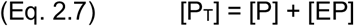

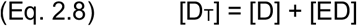

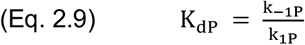

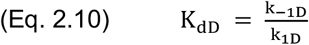

For Eq. 3.1-4.2, apply Eq. 2 notation. Eq. 4 gives relative rate of change in protein-prey complex under initial conditions. Eq. 4.1 allows EP to decay by both dissociation and direct transfer, while Eq. 4.2 represents the classical model with decay by dissociation only. See Theoretical Background for conditions of simplification.

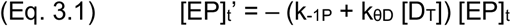

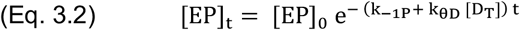

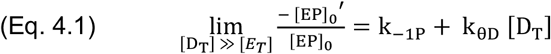

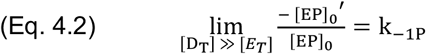

For Eq. 5.1-2, apply Eq. 4 notation, and α and β are arbitrary tuning parameters. See Theoretical Background for conditions of simplification.

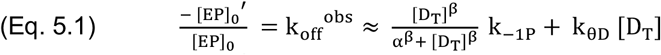

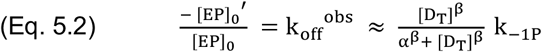

For Eq. 6.1-3, apply Eq. 2 notation, a_1_ and a_2_ are arbitrary constants, and δ is the relative dynamicity of a protein-polynucleotide interaction. See Theoretical Background for conditions of simplification.

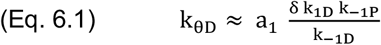

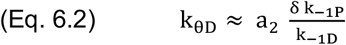

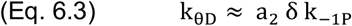

For Eq. 7, FP_E_ is polarization at a given [E_T_], FP_max_ is the maximum polarization, FP_min_ is the minimum polarization, [E_T_] is the total protein concentration, and K_dapp_ is the apparent dissociation constant.

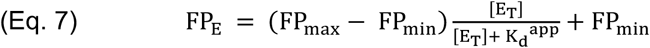

For Eq. 8.1-2, N_t_ is relative polarization at a given time (t), N_min_ is the minimum relative polarization, λ is the exponential rate constant, and k_off_^obs^ is the apparent dissociation rate.

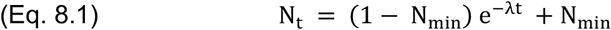

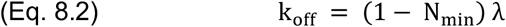

For Eq. 9, apply Eq. 7 notation, [P_T_] is total prey concentration, and S is ligand-protein stoichiometry.

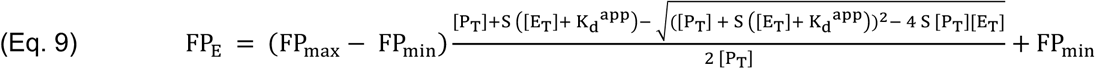

For Eq. 10.1-5, apply Eq. 2 notation.

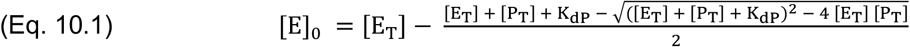

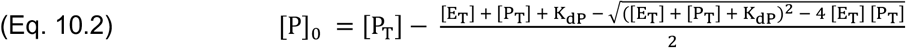

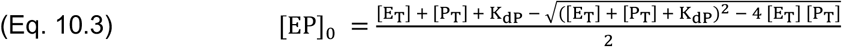

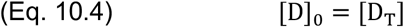

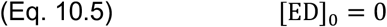

For Eq. 11.1-5, apply Eq. 2 notation, and L1 & L2 are ligand labeled with fluorophores 1 & 2, respectively.

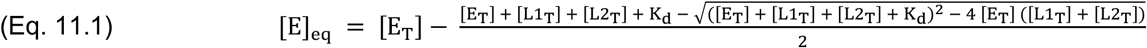

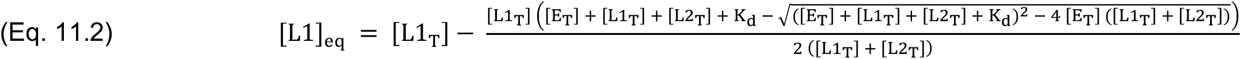

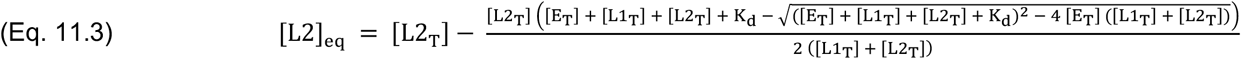

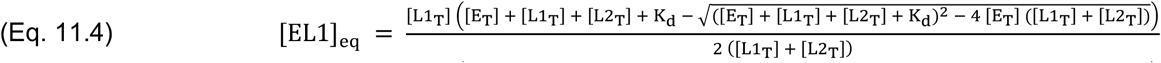

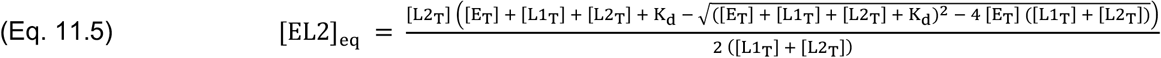

For Eq. 12.1-5, apply Eq. 11 notation, and λ_a→b_ is the rate parameter for an exponential cumulative probability distribution that describes the probability of one particle transitioning from binding state a to b within a given time.

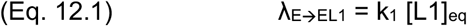

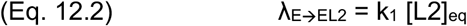

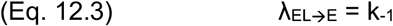

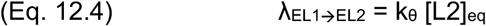

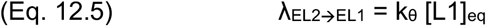

## ACKNOWLEDGEMENTS

W.O.H. was supported by the National Institutes of Health (F32-GM147934). T.R.C. is an investigator of the Howard Hughes Medical Institute.

We’d like to thank Otto Kletzien (Batey lab, University of Colorado Boulder) for the generous gift of recombinant hnRNP-U protein, and Chen Qiu (Hall lab, NIEHS) for the generous gift of FBF-2 protein. We also thank Olke Uhlenbeck, Halley Steiner, Deborah Wuttke, and other members of the Cech lab (University of Colorado Boulder), for stimulating discussion and feedback concerning these studies.

---

**Supplemental Table 1.** Rate Constant Approximation Strategy is Accurate Across Various Simulated Conditions. Reactions were simulated for the simplified reaction scheme (Fig. 1c) as described. Time-point sampling, replication, and Gaussian error were incorporated into each simulated reaction, the erred data for each reaction were regressed with an exponential dissociation curve to determine k_off_^obs^, plots of k_off_^obs^ versus [D_T_] were regressed with Eq. 5 to determine rate constant values, and then Eq. 5.2 and 5.2 were compared with the Bayesian Information Criterion (BIC) (47) to classify the reactions as classic competition or direct transfer. Values reported include the number of simulated data sets that were used (n), the predictive accuracy of the BIC (BIC_acc_), and several previously defined rate constants (Eq. 2 & Fig. 1c – Simplified Reaction Scheme). Reported values from analysis (Analysis) of the simulated (Simulation) data sets are given as mean ± SD across the indicated number of data set replicates (n). Footnotes in the first column apply to the indicated simulation data set (whole row).

Accuracy Metrics for Rate Constant Determination Strategy
| n | K<sub>dP</sub> (nM) | K<sub>dD</sub> (nM) | k<sub>-1P</sub> (s<sup>-1</sup>) |  | k<sub>-1D</sub> (s<sup>-1</sup>) | k<sub>0P</sub> (M<sup>-1</sup>s<sup>-1</sup>) | k<sub>0D</sub> (M<sup>-1</sup>s<sup>-1</sup>) |  | BIC<sub>acc</sub> (%) |
| --- | --- | --- | --- | --- | --- | --- | --- | --- | --- |
|  | Simulation | Simulation | Simulation | Analysis | Simulation | Simulation | Simulation | Analysis | Analysis |
| <sup>a</sup> 20 | 1.0 | 1.0 | 1.0x10<sup>-2</sup> | 1.0 ± 0.025 (x10<sup>-2</sup>) | 1.0x10<sup>-2</sup> | 0 | 0 | -33 ± 46 | 95 (75-100) |
| 12 | 100 | 100 | 1.0x10<sup>-2</sup> | 0.97 ± 0.11 (x10<sup>-2</sup>) | 1.0x10<sup>-2</sup> | 0 | 0 | 75 ± 98 | 92 (62-100) |
| <sup>a,b,c</sup> 19 | 1.0 | 1.0 | 1.0x10<sup>-4</sup> | 0.87 ± 0.079 (x10<sup>-4</sup>) | 1.0x10<sup>-4</sup> | 0 | 0 | -0.76 ± 0.40 | 100 (82-100) |
| <sup>b,c</sup> 16 | 100 | 100 | 1.0x10<sup>-4</sup> | 0.68 ± 0.72 (x10<sup>-4</sup>) | 1.0x10<sup>-4</sup> | 0 | 0 | -5.8 ± 3.3 | 6 (0.15-30) |
| <sup>†</sup> 100 | 10 | 10 | 1.0x10<sup>-3</sup> | 0.99 ± 0.021 (x10<sup>-3</sup>) | 1.0x10<sup>-3</sup> | 0 | 0 | 0.93 ± 2.3 | 97 (91-99) |
| <sup>†</sup> 100 | 10 | 10 | 1.0x10<sup>-3</sup> | 0.99 ± 0.023 (x10<sup>-3</sup>) | 1.0x10<sup>-3</sup> | 100 | 100 | 100 ± 4.0 | 100 (96-100) |
| <sup>†</sup> 100 | 10 | 10 | 1.0x10<sup>-3</sup> | 0.96 ± 0.048 (x10<sup>-3</sup>) | 1.0x10<sup>-3</sup> | 500 | 500 | 500 ± 18 | 100 (96-100) |
| 20 | 10 | 10 | 1.0x10<sup>-3</sup> | 0.97 ± 0.068 (x10<sup>-3</sup>) | 1.0x10<sup>-3</sup> | 0 | 500 | 500 ± 22 | 100 (83-100) |
| 20 | 10 | 10 | 1.0x10<sup>-3</sup> | 1.0 ± 0.021 (x10<sup>-3</sup>) | 1.0x10<sup>-3</sup> | 500 | 0 | 0.76 ± 2.1 | 95 (75-100) |
| <sup>a,c</sup> 11 | 1 | 100 | 1.0x10<sup>-3</sup> | 1.2 ± 0.24 (x10<sup>-3</sup>) | 1.0x10<sup>-3</sup> | 0 | 0 | -12 ± 15 | 45 (17-77) |
| 20 | 100 | 1 | 1.0x10<sup>-3</sup> | 1.0 ± 0.019 (x10<sup>-3</sup>) | 1.0x10<sup>-3</sup> | 0 | 0 | -3.8 ± 2.3 | 95 (75-100) |
| <sup>a,c</sup> 6 | 1 | 100 | 1.0x10<sup>-3</sup> | 1.1 ± 0.13 (x10<sup>-3</sup>) | 1.0x10<sup>-3</sup> | 100 | 100 | 86 ± 13 | 100 (54-100) |
| 19 | 100 | 1 | 1.0x10<sup>-3</sup> | 1.0 ± 0.043 (x10<sup>-3</sup>) | 1.0x10<sup>-3</sup> | 100 | 100 | 98 ± 5.6 | 100 (82-100) |
| 20 | 10 | 10 | 1.0x10<sup>-2</sup> | 1.1 ± 0.33 (x10<sup>-2</sup>) | 1.0x10<sup>-4</sup> | 0 | 0 | -69 ± 220 | 45 (23-68) |
| <sup>b,c</sup> 20 | 10 | 10 | 1.0x10<sup>-4</sup> | 0.83 ± 0.10 (x10<sup>-4</sup>) | 1.0x10<sup>-2</sup> | 0 | 0 | -0.084 ± 0.10 | 100 (83-100) |
| 20 | 10 | 10 | 1.0x10<sup>-2</sup> | 1.1 ± 0.40 (x10<sup>-2</sup>) | 1.0x10<sup>-4</sup> | 100 | 100 | 88 ± 250 | 90 (68-99) |
| 20 | 10 | 10 | 1.0x10<sup>-4</sup> | 0.92 ± 0.079 (x10<sup>-4</sup>) | 1.0x10<sup>-2</sup> | 100 | 100 | 100 ± 1.9 | 100 (83-100) |
| <sup>†,d</sup> 72 | 10 | 10 | 1.0x10<sup>-3</sup> | 0.97 ± 0.090 (x10<sup>-3</sup>) | 1.0x10<sup>-3</sup> | 0 | 0 | 2.8 ± 7.2 | 96 (88-99) |
| <sup>†,d</sup> 44 | 10 | 10 | 1.0x10<sup>-3</sup> | 1.0 ± 0.11 (x10<sup>-3</sup>) | 1.0x10<sup>-3</sup> | 100 | 100 | 100 ± 17 | 91 (78-97) |
| <sup>†,d</sup> 45 | 10 | 10 | 1.0x10<sup>-3</sup> | 2.1 ± 5.8 (x10<sup>-3</sup>) | 1.0x10<sup>-3</sup> | 500 | 500 | 470 ± 93 | 87 (73-95) |

<sup>a</sup> Simulations used [E<sub>T</sub>] = 50 nM; due to K<sub>dP</sub> < [P<sub>T</sub>].
<sup>b</sup> Simulations used 10-hour reaction window; due to low k<sub>off</sub><sup>obs</sup>.
<sup>c</sup> Analyses used simulation-provided baseline signal for k<sub>off</sub><sup>obs</sup> calculations; due to incomplete competition.
<sup>d</sup> Simulations used 20% measurement error.
<sup>†</sup> Simulations generated 100 data sets for analysis.

**Supplemental Table 2.**
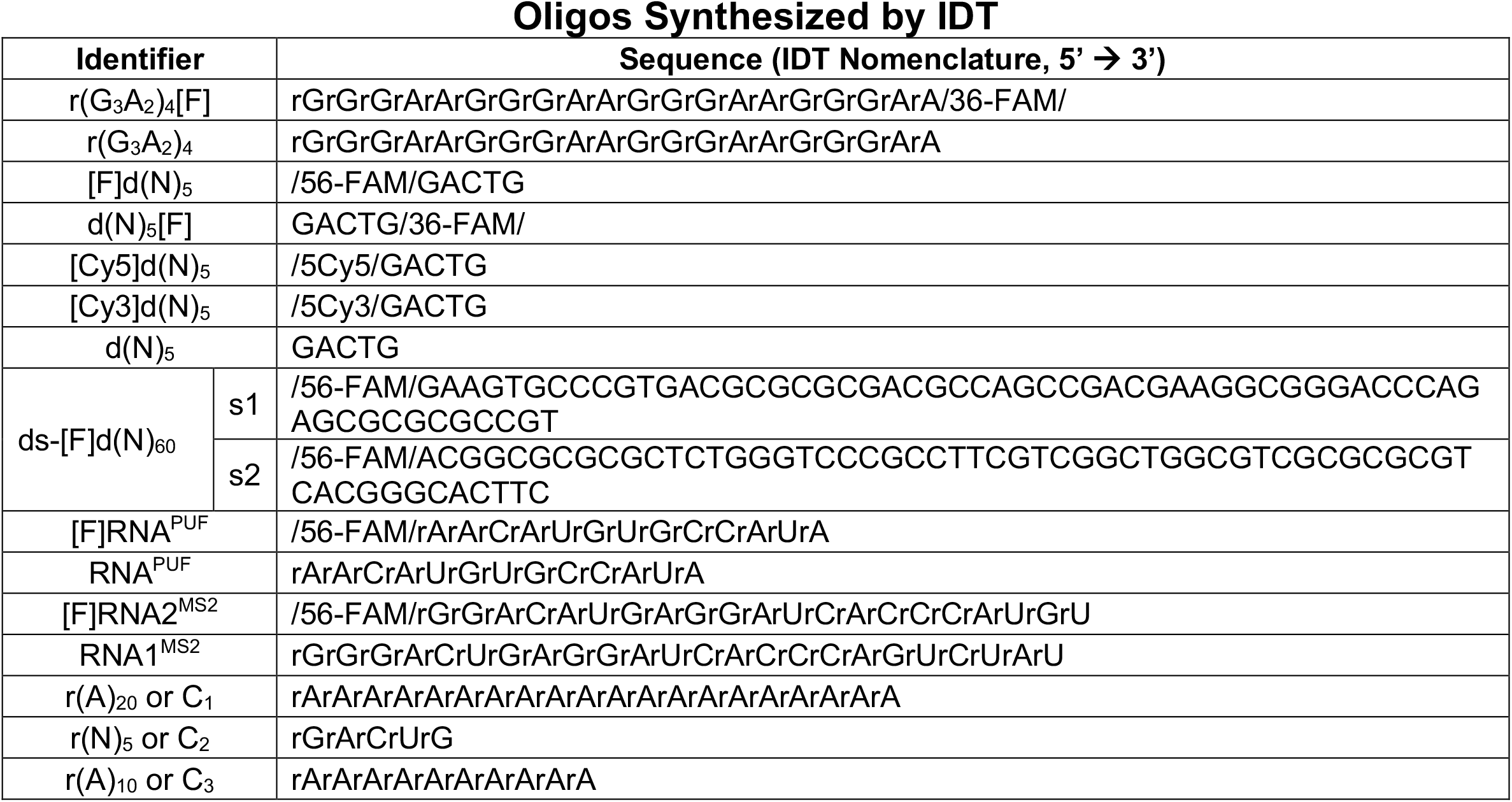
Identities of IDT-Synthesized Polynucleotide Species. The nomenclature used for ordering oligos from IDT (Sequence) is provided for all oligos named (Identifier) in the paper.

**Supplemental Figure 1.**
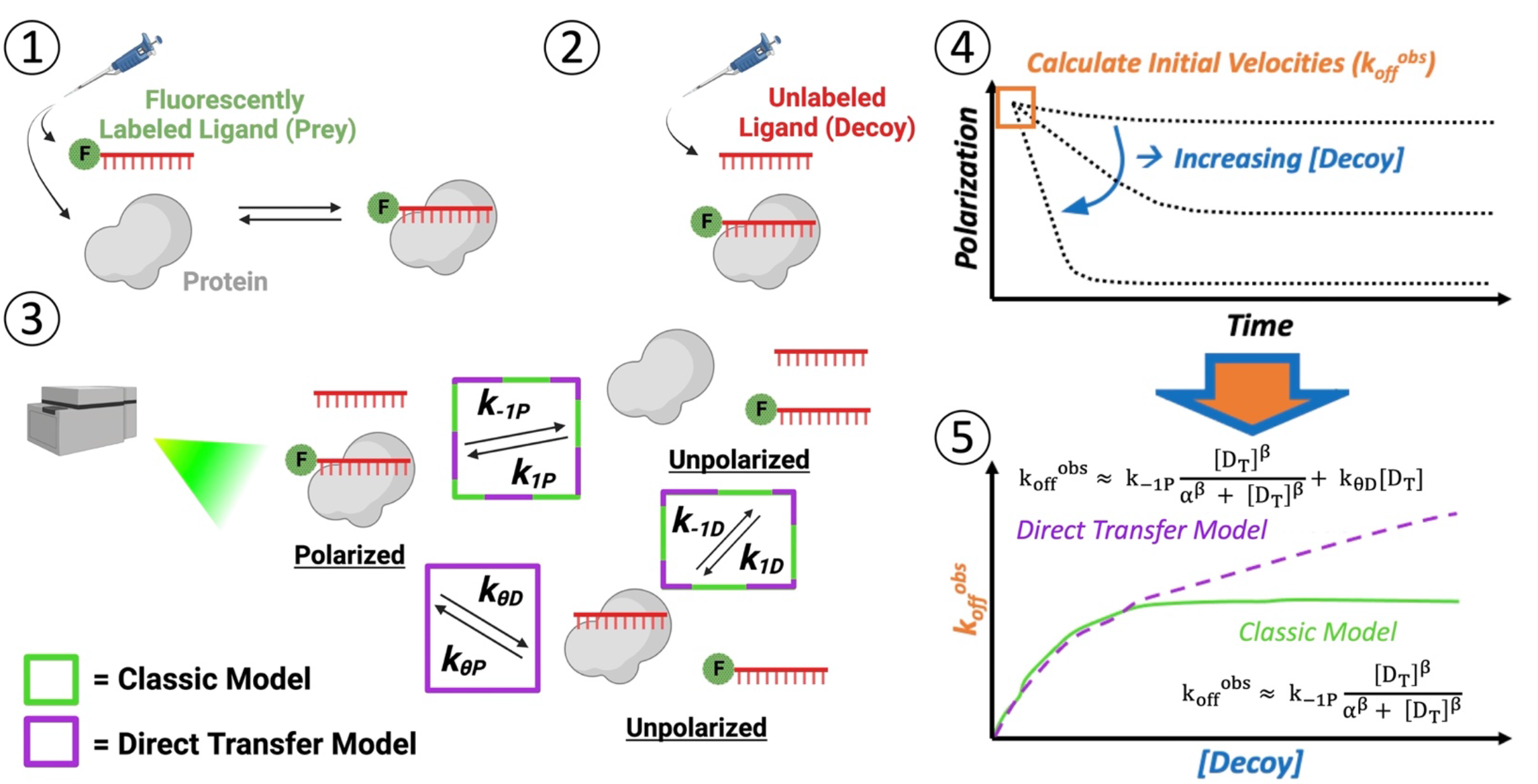
Experimental Strategy to Measure Direct Transfer Kinetics. (1) The minimum amount of protein required for saturated binding is mixed with a trace amount of fluorescently labeled oligonucleotide, then incubated (at 4/25/37 °C) until thermal and reaction equilibrium. (2) Various concentrations of unlabeled decoy oligonucleotide are added to the preformed complex to initiate reactions (at 25°C). (3) The time-course reactions are immediately monitored by fluorescence polarization in a microplate reader (at 25°C). Potential complexes with their polarization states are shown, and they are labeled with rate constants describing inter-complex transitions. Rate constants associated with a classic competition model are indicated by green boxes, and those additionally necessary for a direct transfer model are indicated by a purple box. (4) Polarization signals are normalized to the range in polarization signal across all decoy concentrations to give proportion of initial complex remaining. Normalized polarization signals are plotted versus time and fit with one-phase exponential decay regression. (5) The initial slopes of the regressions (k_off_^obs^) are plotted versus decoy concentration and regressed with custom equations describing the classic competition and direct transfer models to determine rate constant values.

**Supplemental Figure 2.**
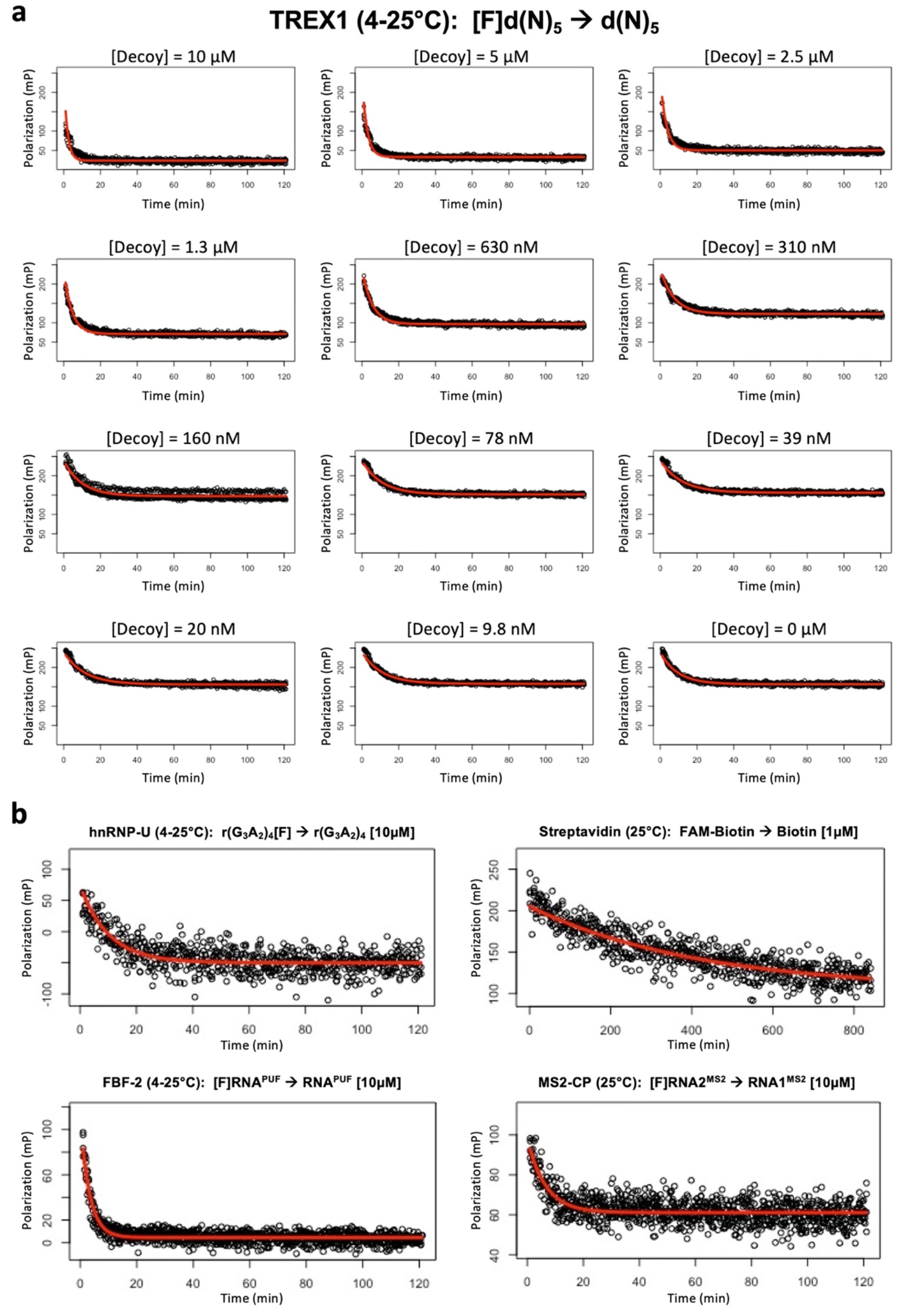
Interrogated Protein-Polynucleotide Interactions Exhibit Pseudo-Exponential Dissociation. **[a]** *A Representative Protein-Ligand Interaction Exhibits Pseudo-Exponential Dissociation Across All Decoy Concentrations*. Raw data (black circles) fit with an exponential decay curve (red line) is shown for all decoy concentrations used in the corresponding experiment in Fig. 2. **[b]** *All Tested Protein-Ligand Interactions Exhibit Pseudo-Exponential Dissociation at the Excess Decoy Concentrations*. Raw data (black circles) fit with an exponential decay curve (red line) are shown for the highest decoy concentrations used in all corresponding experiments in Fig. 2.

**Supplemental Figure 3.**
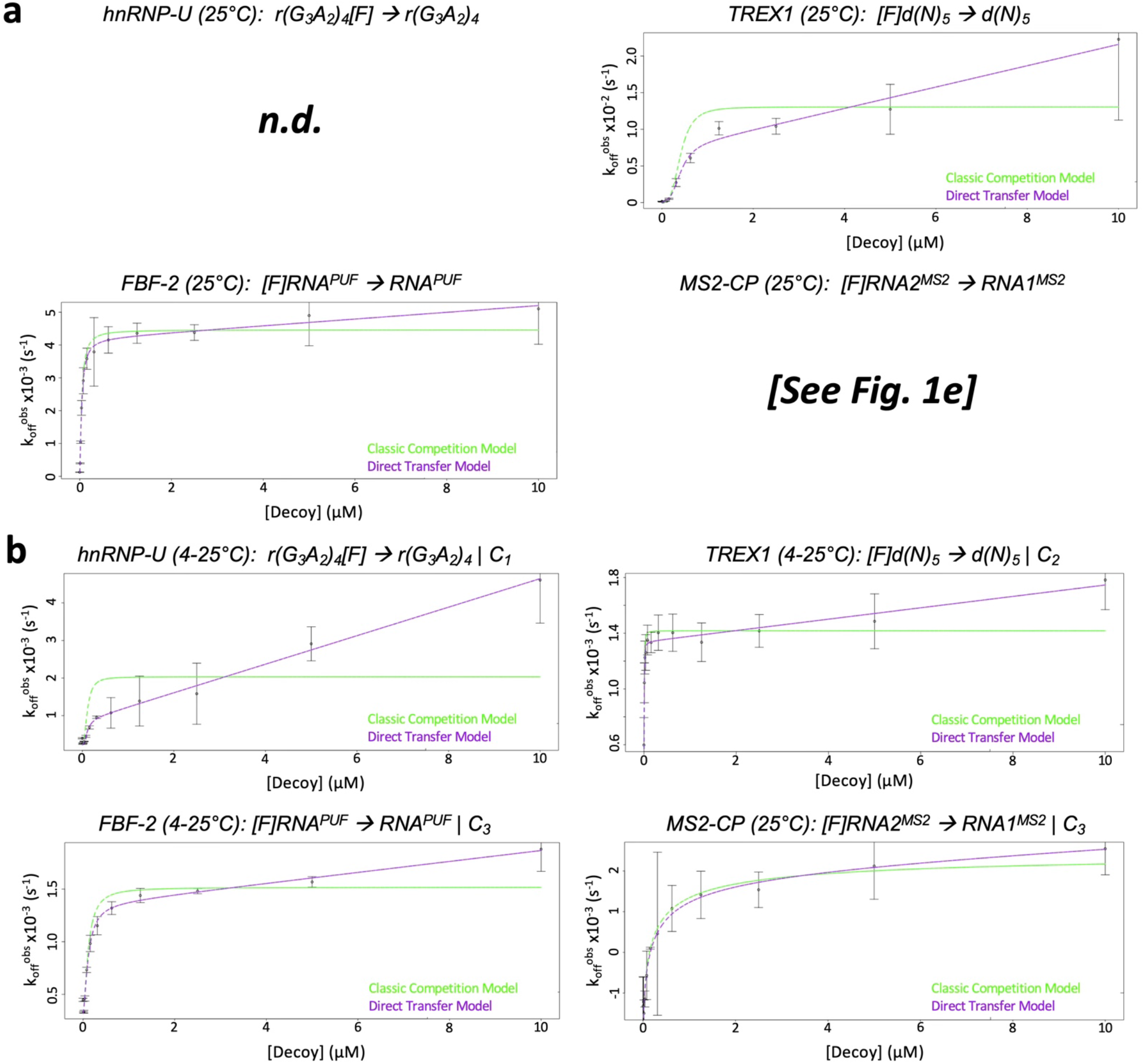
Apparent Direct Transfer is Not an Artifact of Temperature or Polynucleotide Concentration. For all protein-polynucleotide interactions in Fig. 1, fluorescence polarization-based competition experiments were performed and analyzed as described (Supp. Fig. 1) with temperature (panel a) and/or total polynucleotide concentration (panel b) kept constant, and the final plots of apparent prey dissociation rate (k_off_^obs^) versus decoy concentration are shown. Plots show best-fit regression of the data with equations describing classic competition (green lines) versus direct transfer (purple lines). Graphs are from representative experiments, and error bars are mean ± SD across four technical replicates. The corresponding regression values can be found in Table 1.

**Supplemental Figure 4.**
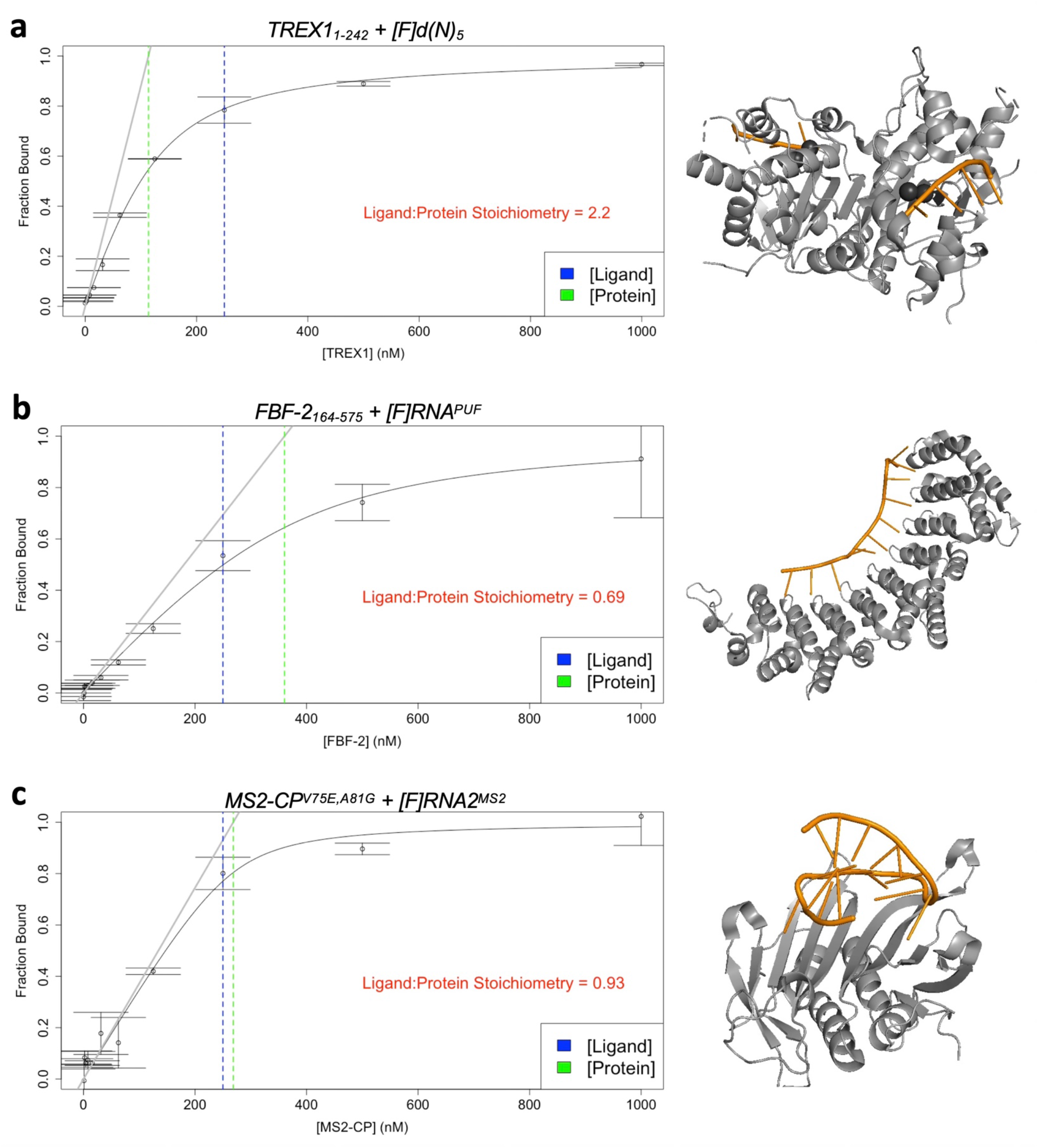
Key Protein-Polynucleotide Interactions Exhibit Expected Stoichiometry. For key protein-polynucleotide interactions, stoichiometry experiments were performed and analyzed as described, and the plots of fraction ligand bound versus protein concentration are shown alongside crystal structures of their interactions. The concentrations of protein (green) and ligand (blue) present at stoichiometric binding equilibrium are shown as vertical dashed lines. The number of ligand molecules inferred to bind to the presumed functional unit of each protein is written in red, and the value should correspond to the number of ligand molecules shown in the corresponding crystal structures. Protein concentrations are for TREX1 homodimer, FBF-2 protomer, and MS2-CP homodimer. Graphs are from single experiments, and error bars are mean ± SD across four technical replicates. Crystal structures show proteins as grey cartoons and ligands as orange cartoons/sticks. Structures in panels a-c are PDB 2OA8, 3V74, and 2C51, respectively.

**Supplemental Figure 5.**
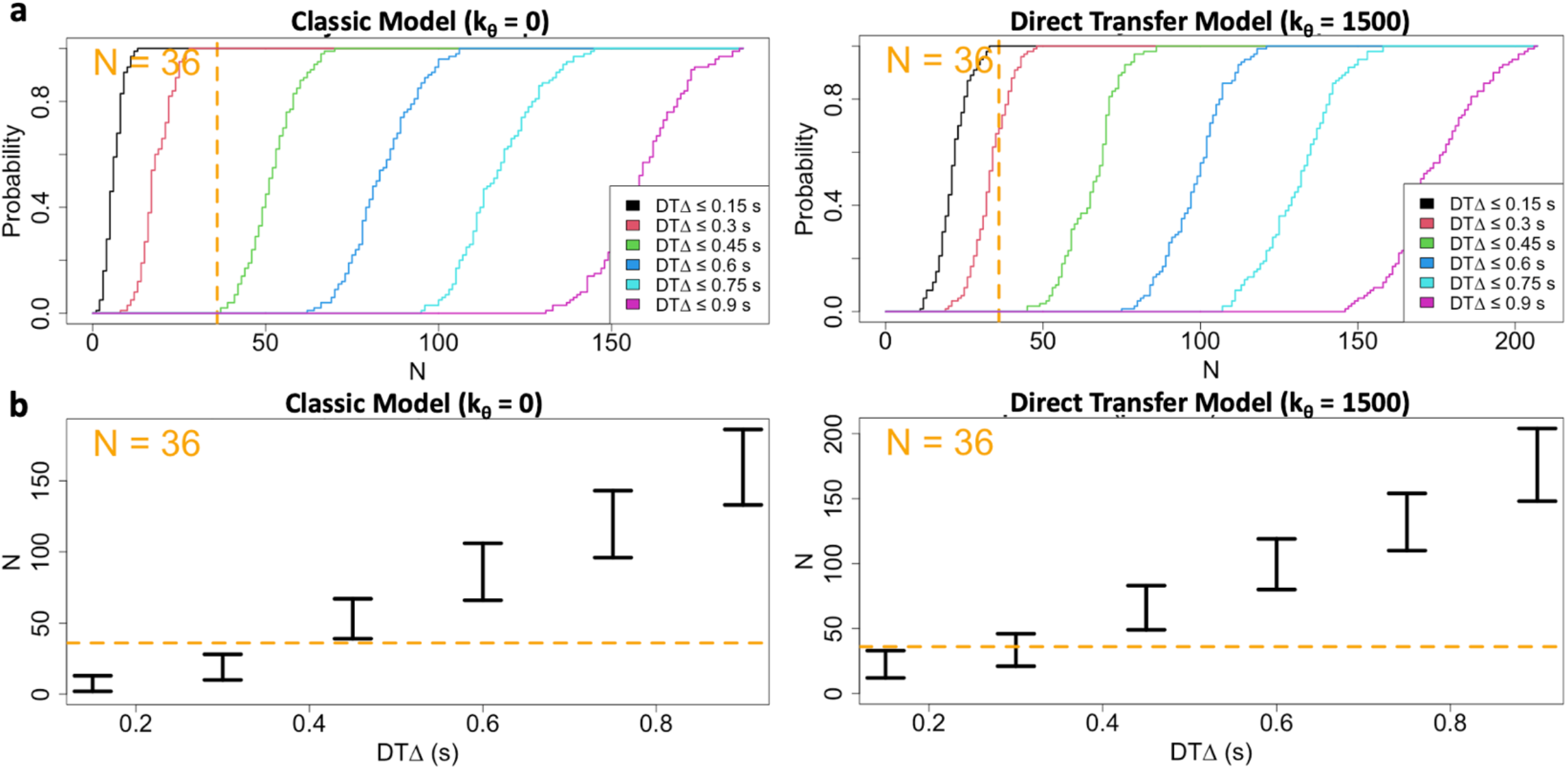
Frequency of Exchange Events in Single-Molecule Data is Consistent with Direct Transfer Model. Dual-label single-molecule experiments (Fig. 2d) were simulated under our experimental conditions (see Methods) using the rate constants reported in Fig. 2c to determine the probable numbers of apparent exchange (i.e., direct transfer) events (N) given thresholds for the apparent time between anti-correlated binding state changes (DTΔ). Panel a shows the cumulative probability distributions of expected N at various DTΔ for the classic and direct transfer models. Panel b shows the 95% confidence intervals of expected N at various DTΔ for the classic and direct transfer models. In all graphs, the dashed orange lines represent our experimentally observed number of apparent exchange events (N=36).

